# Partial EMT Drives Persistent Collective Migration via Collision Guidance in Heterogeneous Populations

**DOI:** 10.64898/2026.04.07.714519

**Authors:** Hyuntae Jeong, Jiwon Kim, Jea-Yun Sim, Susan E. Leggett, Ian Y. Wong

## Abstract

The epithelial-mesenchymal transition (EMT) alters cell-cell interactions to facilitate collective or individual migration during embryonic development, wound repair, or tumor invasion. Epithelial cells are typically cohesive and stationary while mesenchymal cells are individually dispersed and motile. Additional “partial” EMT states are thought to occur with distinct adhesive and migratory behaviors, but these functional phenotypes are poorly understood. Here, we show that cells treated with moderate TGF-*β* concentration exhibit collective migration that is fast and directionally persistent despite heterogeneity in epithelial, partial, and mesenchymal states. We find cells coordinate their motility by reorienting in similar directions after transient contacts, a distinct “collision guidance” mechanism that differs from epithelial arrest or mesenchymal repulsion. Moreover, partial EMT cells sustain collision guidance when interacting with epithelial or mesenchymal cells, which otherwise have increased tendency to repel. We corroborate these experimental observations with a computational model using self-propelled interacting particles that align their motion or repel upon contact. Finally, we show that partial EMT enables tissue monolayer fronts to overwhelm and displace monolayers of other cell types after collision. Overall, these results reveal that partial EMT promotes coherent and emergent behaviors that bridge from cell to tissue length scales, with potential implications for shaping epithelial tissue formation, regeneration, or disorganization.

## I. INTRODUCTION

The epithelial-mesenchymal transition (EMT) is increasingly regarded as a multistep process associated with weakened cell-cell adhesions and increasingly pro-migratory phenotypes during embryonic development, wound healing and tumor progression [1, 2]. Classically, epithelial cells halt their motility upon contact with neighboring cells, which maintains an arrested and tightly-packed tissue architecture (i.e. epithelial contact inhibition of locomotion) [3]. In comparison, mesenchymal cells “repel” upon contact and steer their motility in opposite directions (i.e. mesenchymal contact inhibition of locomotion) [4]. There is increasing interest in “partial” EMT states that may exhibit hybrid or intermediate features between epithelial and mesenchymal states [1, 2]. In particular, partial EMT is associated with collective migration [5–7], especially leader cells that are partially adherent to followers while also expressing vimentin with elongated morphologies [8–10]. (Partial) EMT may be induced by exogenous transforming growth factor beta (TGF-*β*), which drives alterations in single cell morphology and biomarker expression, including a decrease in E-cadherin and increase in N-cadherin [11–20]. However, the functional characteristics of partial EMT states and their contributions towards collective migration remain poorly understood [21].

Collective migration is guided by coordinated mechanical interactions between cells [22]. Typically, densely packed monolayers with very strong cell-cell adhesion tend to be arrested (“jammed”), while more sparse cells with weak cell-cell adhesions are minimally coordinated [23]. Thus, in order for coherent motion to occur, monolayers ought to exhibit moderate density and some intermediate strength of cell-cell adhesion [24]. Such transitions between arrested, individual, and collective motility are often described using minimal physical models assuming identical interacting particles [25–28]. However, this premise is confounded by direct experimental measurements, revealing profound single cell heterogeneity in cell shape [29–31], along with striking fluctuations in cell-cell adhesion [32, 33]. Moreover, heterogeneous populations prepared by co-culturing epithelial and mesenchymal cells are less arrested [34], but can self-sort via differential adhesion [35]. Thus, some population heterogeneity with respect to cell-cell adhesion may facilitate coherent motion in epithelial monolayers, but excess heterogeneity much may impair mechanical coordination across distinct cell states, disrupting collective migration over larger length scales. An unresolved question is how robustly collective migration can occur when EMT promotes heterogeneity in single cell adhesion and motility. This has particular relevance for the formation of interfaces between distinct tissue compartments [36], which has been studied based on collisions between expanding monolayers of different types [37–39].

In this article, we show that epithelial cells treated with moderate TGF-*β* concentrations exhibit fast and directionally persistent collective migration which differs from arrested epithelial cells and uncoordinated mesenchymal cells. Single cell analysis of a dual-color fluorescent EMT reporter revealed that moderate TGF-*β* treatment resulted in the most heterogeneous composition of epithelial (E-cadherin+), partial (E-cadherin-, ZEB1-) and mesenchymal (ZEB1+) cell states. We find that epithelial and partial EMT cells often collide and realign along similar migration trajectories, whereas mesenchymal cells remain dispersed due to uninhibited or repulsive interactions. We simulated these cell-cell interactions using a minimal self-propelled particle model that recapitulates our experimental observations of persistent collective motion. Finally, expanding epithelial monolayers treated with moderate TGF-*β* concentrations exhibit collective fronts that overwhelm epithelial or mesenchymal monolayers upon collision. Thus, cells with partial EMT states exhibit cell-cell interactions distinct from classical epithelial or mesenchymal phenotypes that enable persistent coordinated motility within heterogeneous populations, with potential implications for understanding collective transitions in development and disease.

## II. RESULTS

### A. Epithelial monolayers exhibit coordinated and persistent motility after treatment with moderate but not high TGF-*β*1 concentration

We measured cell migration in monolayers of mammary epithelial cells (MCF-10A) after treatment with 1 ng/mL or 5 ng/mL TGF-*β*1 relative to untreated control. Briefly, MCF-10A cells were cultured for 7 days in a tissue culture flask (either untreated or TGF-*β*1 treated), then labeled with fluorescent stains for the cytoplasm and nucleus, respectively. Cells were replated at 60% confluency onto fibronectin-coated glass bottom plates and allowed to adhere for 24 h, then imaged using fluorescence microscopy under environmentally controlled conditions (Fig. 1a). The velocity fields of the cell monolayers were extracted from time-lapse images using optical flow (Fig. 1b). The spatial coordination in cell velocity was quantified using a correlation length, which measures the characteristic scale over which velocities become uncorrelated (Fig. 1c).

**FIG. 1.**
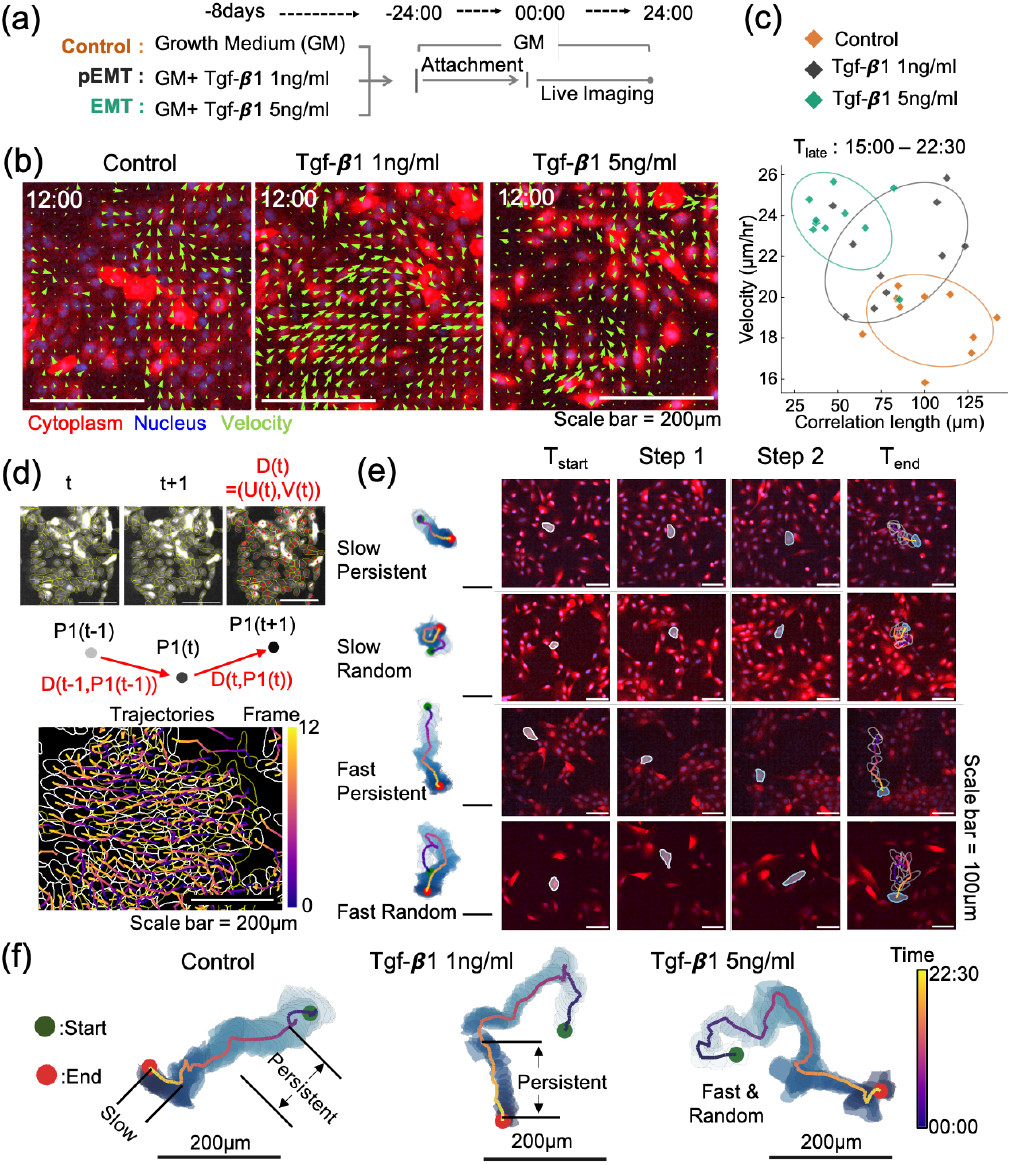
MCF-10A cells after 1 ng/mL TGF-*β*1 treatment exhibit coordinated and faster migration at later times than untreated controls or 5 ng/mL TGF-*β*1 treatment. (a) Experimental design compares cells in untreated control with 7 day treatment with 1 ng ml^−1^ or 5 ng ml^−1^ TGF-*β*1. Cells were then replated for live imaging. (b) Representative snapshots of cell velocity fields for all three conditions (green arrows). Cell cytoplasm is labeled in red (CellTracker Deep Red) and nuclei in blue (Hoechst). (c) Comparison of average cell velocity and correlation length for each experimental condition from 15-22.5 h. Elliptical contours illustrate the Gaussian summaries of the data distribution, constructed from the mean and covariance matrix. (d) Schematic and representative images for single cell trajectories reconstructed from optical flow. (e) Representative cell trajectories classified as slow and persistent, slow and random, fast and persistent, or fast and random. (f) Comparison of representative single cell trajectories for control relative to TGF-*β*1 treatments, revealing distinct behaviors at early and later times.

From 0 to 7.5 h, cells in all three conditions exhibited comparable migration speeds (26 *µ*m/h), but were more coordinated (i.e. larger correlation length) in the untreated condition (Fig. S1a,b,c). However, at later times, cells in the untreated condition exhibited relatively slow motility (∼ 19 *µ*m/h) that was highly correlated (∼ 100 *µ*m) (Fig. 1b,c, Fig. S1d,e). In comparison, cells in the 5 ng/mL TGF-*β*1 condition exhibited faster velocities (∼ 24 *µ*m/h) with minimal correlation (∼ 40 *µ*m) (Fig. S1c,e). Nevertheless, cells from the 1 ng/mL TGF-*β*1 condition exhibited faster migration velocities (∼ 22 *µ*m/h) while maintaining a correlation length (∼ 80 *µ*m) comparable to that of the control, indicating highly coordinated motility (Fig. 1c, Fig. S1d,e). We corroborated these results based on single cell trajectories extracted using optical flow (Fig. 1d, Fig. S2a) and observed four characteristic migration behaviors: i) slow & persistent, ii) slow & random, iii) fast & persistent, and iv) fast & random (Fig. 1e). For example, cells in the control condition were initially fast and persistent (0-6 h) but slow and random later (12-18 h) (Fig. 1f). In comparison, cells in the 1 ng/mL TGF-*β*1 condition were fast and persistent at later times, whereas cells in the 5 ng/mL TGF-*β*1 condition were fast and random. In general, cells were more confluent (*>* 90%) at later times relative to earlier times (∼ 60%), which would result in more frequent interactions with their neighbors. We thus hypothesized that differences in single cell trajectories were due to differences in how motile cells respond to physical contacts with their neighbors.

We quantitatively classified these migration behaviors by analyzing each trajectory over 7.5 h intervals using 17 motility metrics such as speed, persistence, directional variability, arrest dynamics, trajectory topology, and coordination with neighboring cells (Fig. 2a). To visualize the distribution of single cell behaviors, we used Uniform Manifold Approximation and Projection (UMAP) for dimensionality reduction of these 17 motility metrics from 13,700 single cell trajectories (Fig. 2b). Using Spearman correlation, we found that the UMAP1 axis correlated with decreasing velocity and distance traveled (e.g. speed, Euclidean displacement, cumulative distance, etc.), while the UMAP2 axis correlated with more tortuous trajectories and greater temporal fluctuations in speed (e.g. loop score, directionality, maximum step, acceleration) (Fig. 2c). Since UMAP is a nonlinear dimensionality reduction technique, we also verified these trends using principal component analysis (PCA) (Fig. S3).

**FIG. 2.**
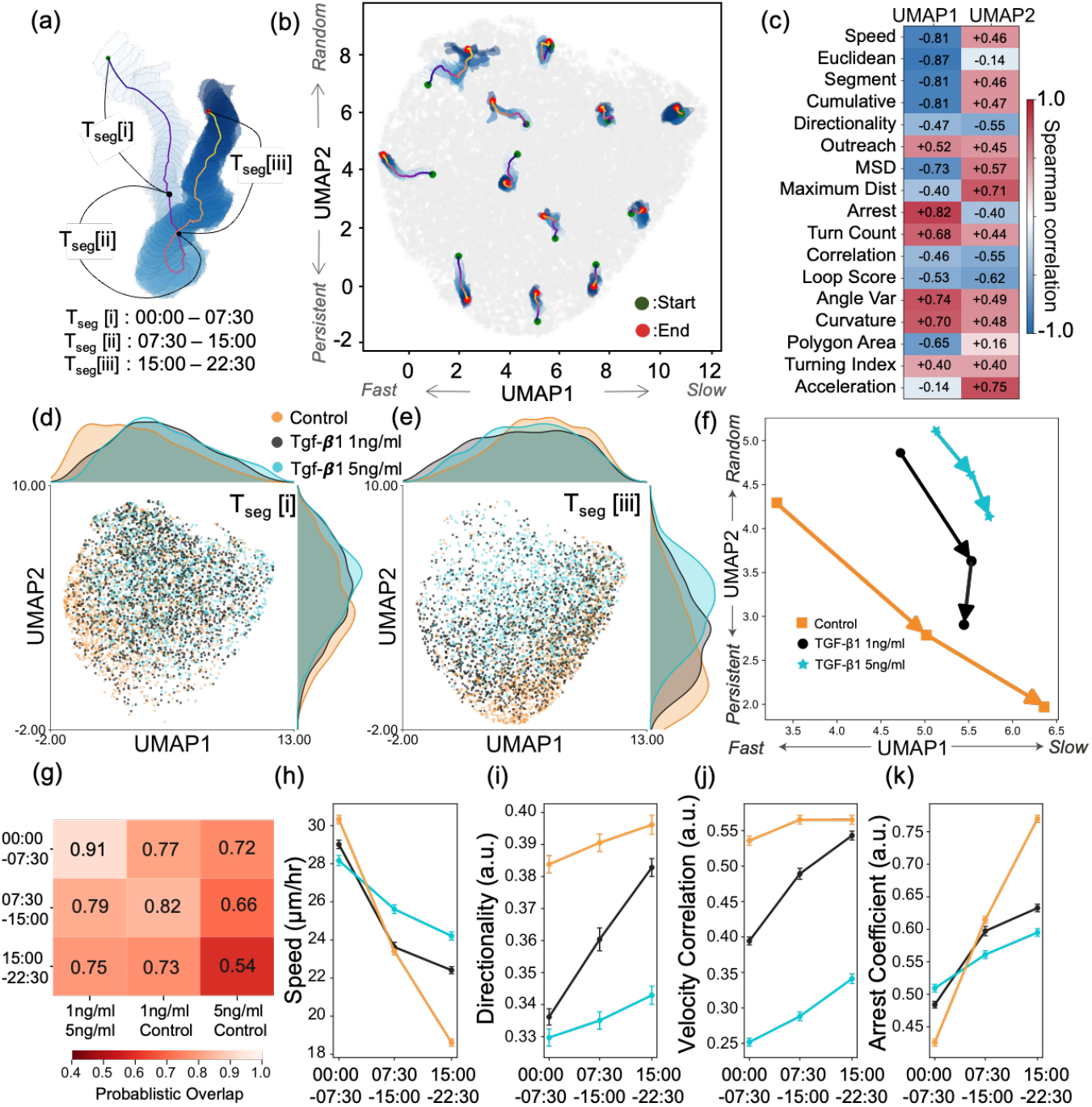
Single cell trajectories after 1 ng/mL TGF-*β*1 treatment are relatively fast and persistent at later times relative to untreated controls or 5 ng/mL TGF-*β*1 treatment. (a) Representative cell trajectory, partitioned into three sequential time segments: *T*_seg_ (i), 00:00–07:30; *T*_seg_ (ii), 07:30–15:00; and *T*_seg_ (iii), 15:00–22:30. (b) UMAP dimensionality reduction of 17 motility metrics for each 7.5 h segment (per trajectory) projected into two dimensions, and overlaid with illustrative single cell trajectory segments. UMAP1 correlates with fast to slow migration, while UMAP2 correlates with persistent to random migration. (c) Spearman correlation between UMAP components and 17 individual motility features. (d, e) Distribution of cell trajectory segment metrics from each experimental condition at time segments *T*_seg_ (i) and *T*_seg_ (iii) (Control, brown; TGF-*β*1 1 ng ml^−1^, gray; and TGF-*β*1 5 ng ml^−1^, mint). Kernel density distributions of each condition along the UMAP1 and UMAP2 components are shown in the top and right panels, respectively. (f) Centroids for each time segment and experimental condition in the UMAP projection. Arrows indicate the displacement of centroids between adjacent segments, revealing changes in motility behavior over time. (g) Pairwise overlap analysis indicating relative similarity between experimental conditions and time windows. Smaller values indicate greater differences in motility behavior. Temporal changes in representative motility features across time segments: (h) speed, (i) directionality, (j) velocity correlation, and (k) arrest coefficient. Each dot represents the mean value of the corresponding motility feature, and error bars indicate the standard error of the mean (SEM).

We found that migration behaviors from all three conditions were comparable at early times (0-7.5 h), although the untreated control distribution was slightly skewed towards lower UMAP1 values, indicating faster motility (Fig. 2d). At later times (15-22.5 h), the untreated and 1 ng/mL TGF-*β*1 distributions were skewed towards higher UMAP1 and lower UMAP2 values relative to early times (0-7.5 h), while the cell distribution of 5 ng/mL TGF-*β*1 remained comparable for early and late times (Fig. 2e). These changes were then quantified based on the position of the centroids of these distributions over the three successive time intervals (Fig. 2f). The untreated distribution showed the most pronounced shift from random and fast at early times to persistent and slow at later times, which we interpret as the onset of collective migration at increased cell density. The 1 ng/mL TGF-*β*1 distribution also exhibited some slowdown at later times, with a more dramatic shift toward increasing persistence. Finally, the 5 ng/mL TGF-*β*1 distribution exhibited a slight slowdown and increase in directional persistence, but these were relatively subtle relative to the untreated and 1 ng/mL TGF-*β*1 distribution (Fig. 2f). Indeed, the overlap of data distribution also showed that motility behaviors were more similar between 1 ng/mL and 5 ng/mL TGF-*β*1 conditions than between either of these and the untreated control at early times (0 - 7.5 h) (Fig. 2g). However, at later times, motility behaviors under the 5 ng/mL TGF-*β*1 condition became more distinct from those under the 1 ng/mL TGF-*β*1 and untreated conditons (Fig. 2g). These condition-dependent centroid shifts in the UMAP embedding were also observed using PCA (Fig. S3ac). We then directly examined the highly correlated motility metrics, including mean speed, directionality, velocity correlation, and arrest coefficient, revealing that the 1 ng/mL TGF-*β*1 condition showed a marked increase in directional persistence and local coordination, accompanied by a modest reduction in speed (Fig. 2h,i,j,k). Thus, cells in the 1 ng/mL TGF-*β*1 condition exhibit a gradual increase in directionally persistent migration without the slowdown observed for cells in the untreated control.

### B. Increased heterogeneity in fluorescent EMT reporter expression after treatment with moderate TGF-*β* concnetration

These differences in cell motility after 1 ng/mL TGF-*β*1 treatment might occur due to a homogeneous change in EMT state across the entire population or more heterogeneous changes across some subpopulations. In order to resolve EMT at the single cell level after TGF-*β*1 treatment, we used a dual color fluorescent reporter for epithelial, partial, or mesenchymal states. The dual-ZCAD fluorescent reporter indicates an epithelial-like state using red fluorescent protein (RFP) tagged on the E-cadherin promoter, and mesenchymal-like state using green fluorescent protein (GFP) signal expressed on the knockdown of miR-200, which drives the EMT transcription factor Zinc Finger E-Box Binding Homeobox 1 (ZEB1) [16]. For MCF-10A cells expressing the dual Z-cad reporter, we classified EMT state based on the ratio of red and green reporter intensities measured within each cell body (Fig. S2b). For simplicity, we denote red fluorescent cells (E-cadherin^+^) as “E,” uncolored cells (E-cadherin^−^ and ZEB1^−^) as “P” and green fluorescent cells (ZEB1^+^) as “M.”

The cell population in the untreated control was ∼60% red (E) with ∼30% uncolored (P), and the remainder green (M) (Fig. 3a,b). In comparison, the population in the 1 ng/mL TGF-*β*1 condition had ∼10% red (E) with ∼50% uncolored (P) and ∼20% green (M). Finally, the population in the 5 ng/mL TGF-*β*1 condition was ∼5% red (E) with ∼40% uncolored (P) and ∼40% green (M). Markov chain analysis revealed some interconversion between red fluorescence and uncolored in the control, interconversion from red and green to uncolored for 1 ng/mL TGF-*β*1 treatment, and interconversion between uncolored and green for 5 ng/mL TGF-*β*1 treatment (Fig. S2c). Cells were further classified by shape, but the distributions of compact (∼ 40%), elongated (∼ 30%) and spread (∼ 30%) morphologies remained comparable for untreated and TGF-*β*1 conditions (Fig.3c; S4).

**FIG. 3.**
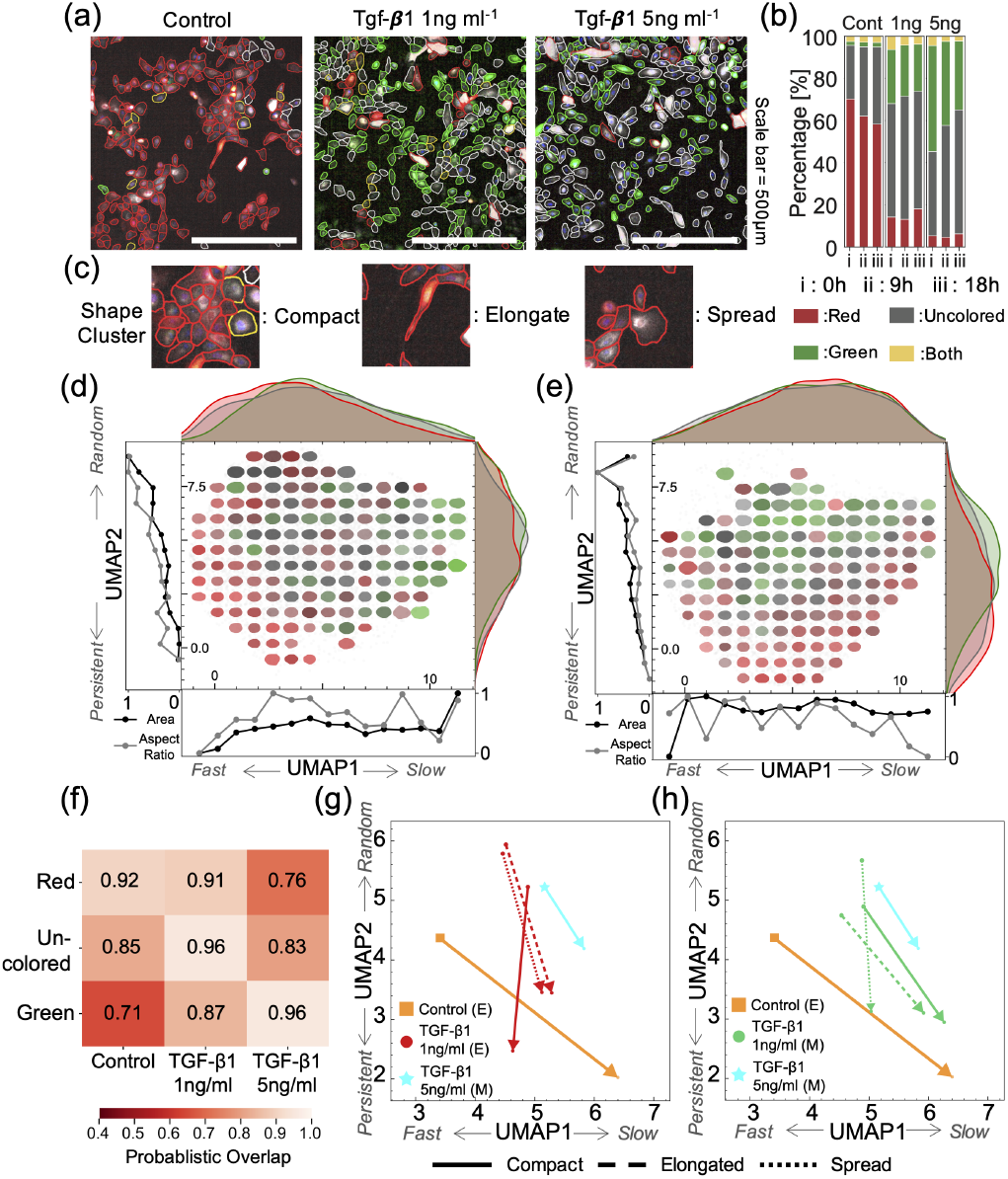
Partial and full EMT states measured using Z-cad dual fluorescent reporter after TGF-*β*1 treatment. (a) Representative images of MCF-10A expressing Z-cad dual fluorescent reporter in untreated control relative to 1 ng/mL and 5 ng/mL TGF-*β*1 treatments. (b) Distribution of EMT reporter color expression at 0, 9, and 18 hours after imaging in control vs TGF-*β*1 treatment. (c) Representative cell morphologies defined as compact, elongated, or spread. (d, e) Distribution of cell color and morphology in the UMAP latent space at different time segments *T*_seg_ (i);(d), *T*_seg_ (iii);(e). Cell shapes are reconstructed from autoencoder-derived latent representations, while color indicates the average EMT reporter level computed over local grid regions in the UMAP space. Subpanels summarize color distributions and representative morphological features. (f) Pairwise overlap analysis indicating relative similarity in motility behavior between EMT reporter and experimental condition. (g, h) Centroids for each time segment for fluorescent reporter state and experimental condition. Arrows indicate the displacement of centroids between adjacent segments, revealing changes in motility behavior over time.

In order to understand how fluorescent reporter expression and morphology map to motility, we projected auto-encoded cell colors and shapes onto our previously generated UMAP representation for cell motility (Fig. 3d,e). We observed that there was a high correlation in UMAP distribution between treatment condition and EMT reporter expression, since the control was mostly red (E), 1 ng/mL TGF-*β*1 was largely uncolored (P), and 5 ng/mL TGF-*β*1 was largely green (M) (Fig. 3a,b). Indeed, the distribution of red (E) and uncolored (P) cells shifted over time from fast to slow (with increasing UMAP1), along with increasing persistence (with decreasing UMAP 2) (Fig. 3d, e). In comparison, green (M) cells maintained similar distributions over time with more random motility (high UMAP2) than red (E) or uncolored (P) cells (Fig. 3d, e). Probabilistic overlap analysis showed that UMAP distributions for control, 1 ng/mL and 5 ng/mL TGF-*β*1 conditions were predominantly associated with the UMAP distributions for red (E), uncolored (P), and green (M), respectively (Fig. 3f). Nevertheless, the 1 ng/mL TGF-*β*1 and uncolored distributions were also similar to the red and green or control and 5 ng/mL TGF-*β*1 conditions, respectively. When considered over sequential time windows, red (E) cells in the control condition exhibited a similar slowdown and increase in persistence as the entire population of control cells over time (Fig.2f; 3g). Green (M) cells in the 5 ng/mL TGF-*β*1 treatment condition also behaved similarly over time, comparable to the entire population of cells in the 5 ng/mL TGF-*β*1 treatment condition (Fig.2f; 3g). However, red (E) cells in the 1 ng/mL TGF-*β*1 treatment condition exhibited shape-dependent differences over time. Elongated and spread cells slowed down slightly with increasing persistence (decreasing UMAP2), while compact cells sped up slightly with increasing persistence (Fig. 3g). In comparison, spread green (M) cells in the 1 ng/mL TGF-*β*1 treatment condition also exhibited increasing persistence over time (decreasing UMAP2), while compact and elongated green (M) cells exhibited some slowdown (Fig. 3h). Altogether, these results show that cell migration behaviors are largely correlated with fluorescent reporter expression for control and 5 ng/mL TGF-*β*1 treatment conditions. However, cell migration behaviors exhibited a more complicated dependence on fluorescent reporter expression for the 1 ng/mL TGF-*β*1 treatment, where some morphological subpopulations exhibited more persistent motion.

### C. Persistent collective migration is robust to heterogeneous EMT states

Although differences in single cell motility correlate with different EMT states, we have not yet addressed the possibility of cooperative behavior between different subpopulations. We analyzed the heterogeneous composition of each population using a ternary phase diagram with three different axes that represent the percentage of cells that are red (E), uncolored (P), or green (M) (Fig. 4a). In this representation, completely homogeneous populations (i.e. 100% of the same color) are located at the corners, while a perfectly heterogeneous population is located at the center (i.e. 33% of cells associated with each color). Thus, cells in the control conditions were mostly red and localized towards the left side of the ternary phase diagram, while cells in the 5 ng/mL TGF-*β*1 conditions were more green and localized towards the right side of the ternary phase diagram. Finally, cells in the 1 ng/mL TGF-*β*1 treatment condition were typically partway between the left and right sides. We quantified the heterogeneity in fluorescent reporter expression based on the relative distance *D* from the origin, normalized by the distance between the origin and any corner *D*_*max*_. There was considerably more heterogeneity in the 1 ng/mL TGF-*β*1 condition relative to the control or 5 ng/mL TGF-*β*1 condition (Fig. 4b).

**FIG. 4.**
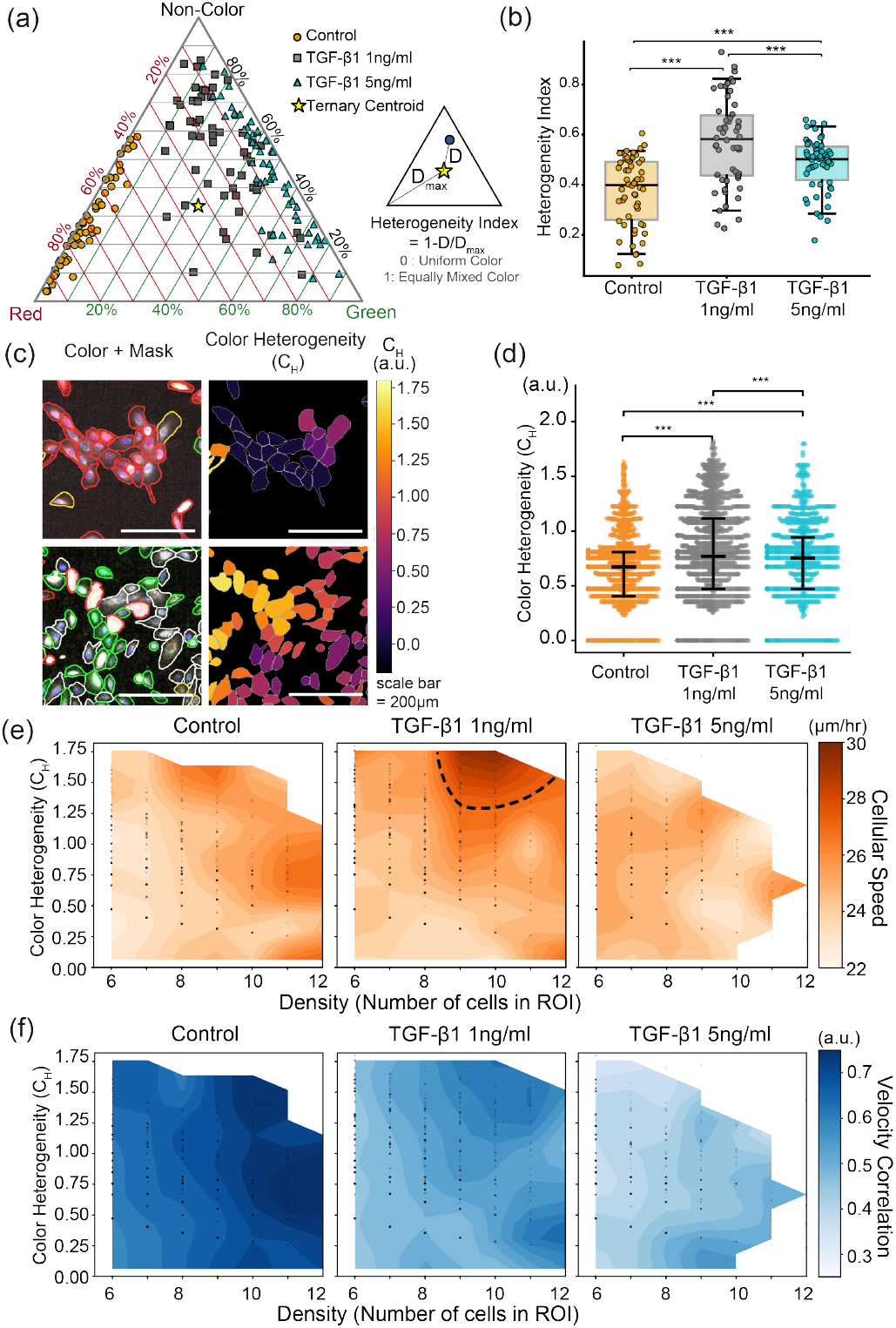
Increased local heterogeneity in Z-cad fluorescent reporter expression after 1 ng/mL TGF-*β*1 treatment relative to control or 5 ng/mL TGF-*β*1 treatment. (a) Ternary diagram showing the percent composition of fluorecent reporter expression(red, green, and uncolored) across experimental conditions. Orange circles: Control; gray squares: TGF-*β*1 (1 ng mL^−1^); cyan triangles: TGF-*β*1 (5 ng mL^−1^). (b) Heterogeneity index is a read out of how evenly the three EMT states are distributed, where 0 corresponds to equal percentages of each EMT state in the population and 1 corresponds a population wholly comprised of one EMT state. Box plots show median and interquartile range with overlaid individual data points. Pairwise comparisons were evaluated using Bonferroni-corrected post hoc tests (*** *p <* 0.001). (c) Representative images of local color heterogeneity within a small region of interest. (d) Color heterogeneity in control or TGF-*β*_1_ treatment condition. Horizontal lines indicate the median and interquartile range (IQR); dots denote individual cells. Statistical significance was assessed using Bonferroni-corrected pairwise comparisons (*** *p <* 0.001). Density–heterogeneity maps of cell motility metrics; (e) speed and (f) velocity correlation. Data were binned along number of cells and color heterogeneity of each patch. Overlaid points represent individual measurements.

We then considered whether this population heterogeneity was also apparent at the single cell level, since different cell types could conceivably self-sort into their respective subpopulations. We quantified local heterogeneity *C*_*H*_ by measuring differences in color expression and cell morphology among neighboring cells within a ∼100 *µ*m wide region of interest of, typically containing 6–12 cells, then sampled across the entire field of view of ∼1000 *µ*m (Fig. S5a, Fig. 4c). This local heterogeneity *C*_*H*_ was significantly higher in 1 ng/mL TGF-*β*1 condition relative to control and 5 ng/mL TGF-*β*1 conditions (Fig. 4d). Moreover, regions with increased cell density and local heterogeneity also exhibited higher average velocity in the 1 ng/mL TGF-*β*1 condition, while remaining comparable in the control and 5 ng/mL TGF-*β*1 conditions (Fig. 4e, S5d,e). For this 1 ng/mL TGF-*β*1 condition, the increased cell density and local heterogeneity was further associated with locally coordinated velocities (Fig. 4e,f). It should be noted that 5 ng/mL TGF-*β*1 treated cells were typically more spread, and so typical regions of interest contained fewer cells than the other conditions, resulting in limited data at high density (shown using white space). Moreover, control and 5 ng/mL TGF-*β*1 treated populations were more homogeneous than 1 ng/mL TGF-*β*1, so there was also limited data for high local heterogeneity *C*_*H*_ (also shown using white space). To corroborate these results, we prepared 1:1 mixtures of untreated and 1 ng/mL or 5 ng/mL TGF-*β*1 treated cells (Fig. S6a,b). Both these mixed populations exhibited enhanced collective motility, characterized by sustained migration speed and persistence at high density and later times (Fig. S6). Overall, coordinated and persistent migration was relatively robust to local heterogeneity in EMT state for moderate 1 ng/mL TGF-*β*1 treatment.

### D. Partial EMT cell states facilitate collision guidance with epithelial and mesenchymal cells

Physically, coordinated and persistent motion in a group of cells occurs through some interaction mechanism where cells match their speed and direction with their neighbors [24]. Visual inspection of single cell trajectories before and after a cell-cell interaction occurred revealed distinct behaviors associated with EMT state (Fig. S7a). For example, “collision guidance” occurred when a cell meeting another cell aligned its direction of motion to match (Fig. 5a) [4]. Alternatively, “repulsion” occurred when a cell meeting another cell would immediately redirect its motion away from the contact (i.e. mesenchymal contact inhibition of locomotion) (Fig. 5b). Finally, “uninhibited” interactions occurred if a cell can migrate over (part of) another cell without changing direction (Fig. 5c). These behaviors were quantified based on the angular change in migration direction Δ*θ*_*reorient*_ for a given cell after contact, and the angle between the migration directions *θ*_*between*_ of the two cells after contact (Fig. 5d). When cells aligned their motility, the resulting angle tended to be lower (*θ*_*between*_ < 80°). We further classified these events with smaller changes in direction as “collision alignment” (Δ*θ*_*reorient*_ < 80°) and larger changes in direction as “collision guidance” (80° ≤ Δ*θ*_*reorient*_). In comparison, “uninhibited” interactions typically resulted in larger angles between cell migration directions (80° ≤ *θ*_*between*_) with smaller changes in direction (Δ*θ*_*reorient*_ < 80°), while “repulsive” interactions showed larger changes in direction (80° ≤ Δ*θ*_*reorient*_).

**FIG. 5.**
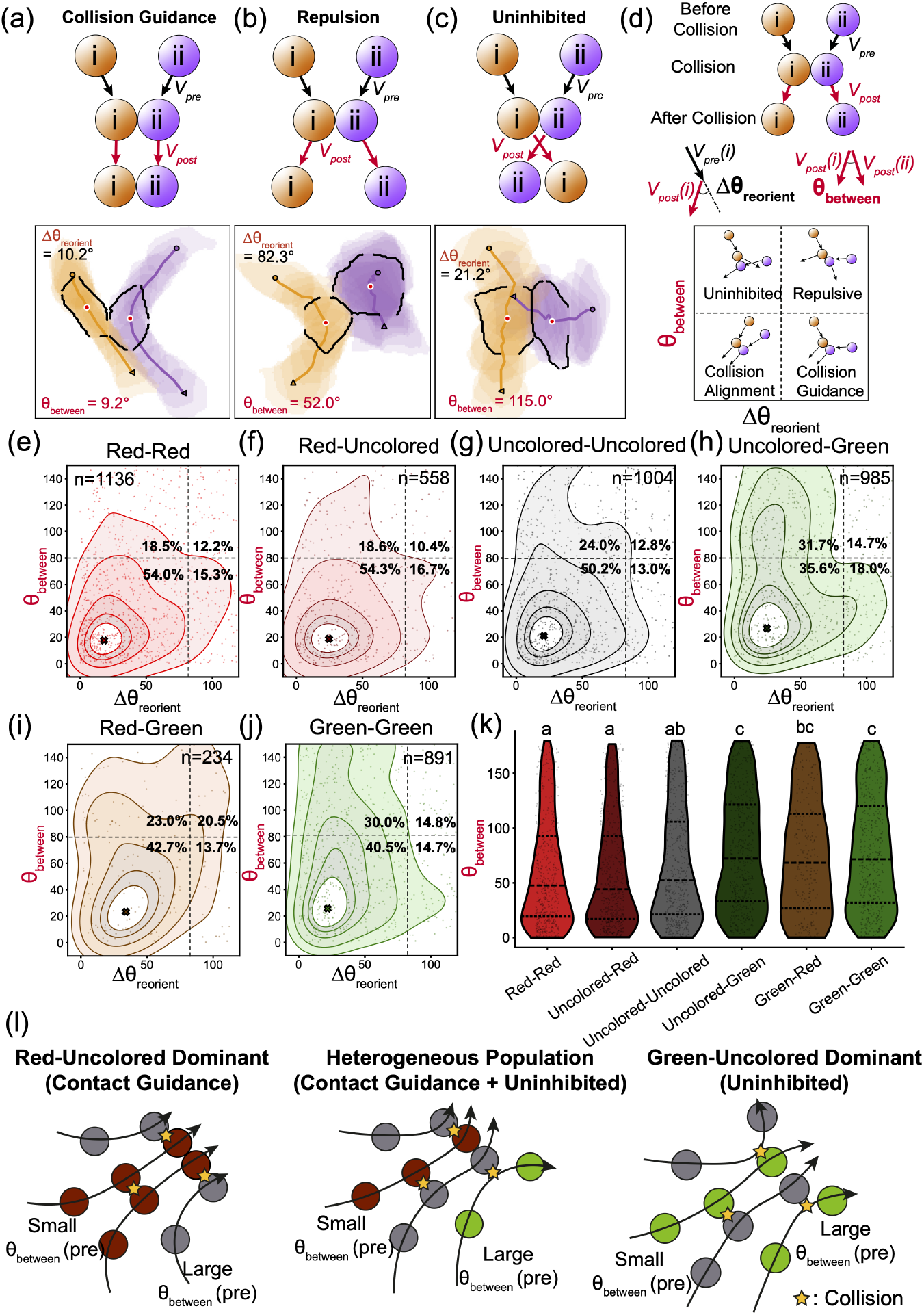
EMT state correlates with distinct collision guidance, uninhibited or repulsive interaction. (a-c) Representative collision scenarios illustrating distinct values of Δ*θ*_reorient_ and *θ*_between_. Low *θ*_between_ corresponds to collision guidance behavior(a), whereas high Δ*θ*_reorient_ indicates strong repulsive reorientation(b). In contrast, low Δ*θ*_reorient_ combined with high *θ*_between_ characterizes uninhibited(sliding) behavior (c). (d) Schematic illustrating the quantitative analysis of cell–cell collision events. Collision responses were quantified by analyzing angular transitions before and after cell–cell contact. Δ*θ*_reorient_ were characterized by the difference between the angle of incidence and the angle of refraction, whereas *θ*_between_ was quantified by measuring the angular difference between the trajectories of interacting cells after collision. 4 different collision modes marked on the coordination space of Δ*θ*_reorient_ and *θ*_between_. (e-j) Kernel density estimate contour lines (level = 0.4, 0.6, 0.8, 0.9,and 0.95) of collision-induced angular responses, plotted as the reorientation angle (Δ*θ*_reorient_) versus the intercellular angle (*θ*_between_) Each figures describe different EMT reporter color pairings ((e)red-red,(f)red-uncolored, (g) uncolored-uncolored, (h)uncolored-green, (i)red-green, and (ij) green-green). (k) Violin plots of *θ*_between_ across collision pairs. Dashed lines present median and interquartile ranges. Statistical differences were assessed using Kruskal-Wallis test followed by pairwise Mann-Whitney U test with Bonferrnoi correction. Group sharing the same letter are not significantly different. (l) Scenarios for the emergence of migration patterns from collision modes in heterogeneous populations.

Interactions between two red fluorescent (E) cells mostly occurred through collision alignment (54%) or collision guidance (15%), whereas uninhibited (19%) or repulsive (12%) were less common (Fig. 5e). The distribution of interaction events between red fluorescent (E) and uncolored (P) cells were comparable with those observed for two red fluorescent (E) cells (Fig. 5e,f). Unexpectedly, interactions between uncolored (P) and green fluorescent (M) cells exhibited increased collision guidance (19%), particularly compared to interactions between two uncolored cells (Fig. 5gh). Further, interactions between red fluorescent (E) and green fluorescent (M) cells exhibited increased repulsion (21%) relative to two uncolored cells, two green fluorescent cells, or an uncolored cell interacting with a green fluorescent cell (Fig. 5g-j). Interactions involving uncolored cells and green fluorescent cells also exhibited more uninhibited events (24-32%) than interactions between two red fluorescent (E) cells (Fig. 5e,h). Thus, uncolored (P) cells exhibited distinct interaction behaviors with red fluorescent (E) cells or green fluorescent (M) cells (respectively) that differed from how red fluorescent cells interact with green fluorescent cells (Fig. 5k). Altogether, red fluorescent (E) and uncolored (P) cells appeared to exhibit collision guidance that sustains collective migration (Fig. 5l). However, the presence of green fluorescent (M) was destabilizing for collective migration by increasing uninhibited and repulsive interactions, particularly with red fluorescent (E) cells. It is intriguing that uncolored (P) cells can sustain collision guidance with green fluorescent (M) cells (at high angles) instead of the repulsion that occurs for red fluorescent (E) cells (Fig. S7). The increased fraction of uncolored (P) cells may be crucial to coordinate both red fluorescent (E) and green fluorescent (M) cells through collision guidance, which would be less effective in a population of only red and green fluorescent cells.

To elucidate these combinatorial interactions within heterogeneous populations, we implemented a minimal computational model to simulate cell-like particles with distinct collision behaviors that recapitulated our experimental observations (Fig. 6ab). Briefly, “epithelial” particles collided inelastically and aligned their velocities after interacting, analogous to the “collision guidance” behavior observed experimentally (Fig. 5f, and Fig. 6c). Instead, “mesenchymal” particles collided elastically and reoriented their subseuqent velocities away from each other. This interaction rule led to reorientation without sustained alignment, consistent with “uninhibited” and “repulsive” interactions observed experimentally (Fig. 5f, and Fig. 6d). A similar interaction occurred when a “mesenchymal” particle interacted with an “epithelial” particle. Particle motion was driven by persistent stochastic forcing via Ornstein-Uhlenbeck acceleration noise and particles could further realign their velocity to match nearby neighbors using a Vicsek-type rule. We initialized 600 particles with uniformly random locations within a circular domain of approximately 30 cell diameters that presented reflecting boundary conditions (Fig. 6b). We again quantified the local color heterogeneity by considering approximately 6-7 neighboring particles around each reference cell (*r* = 3.5) (Fig. S8a). This local color heterogeneity was highest for a 50/50 mix of “epithelial” and “mesenchymal” particles, which were locally well mixed locally based on the standard deviation of local color heterogeneity (Fig. S8a).

**FIG. 6.**
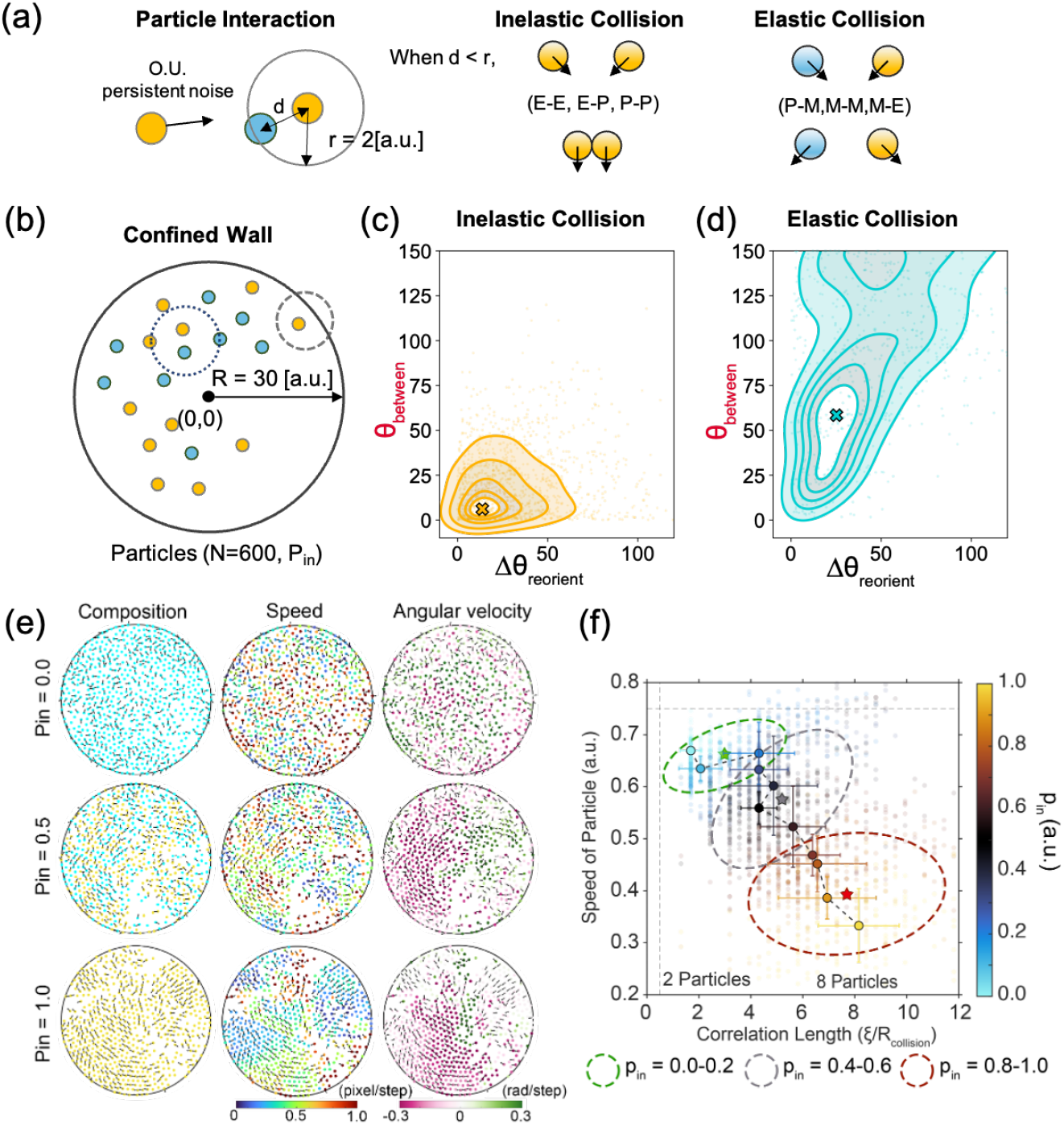
Computational model of coordinated motility in heterogeneous mixtures of self-propelled particles undergoing inelastic or elastic collisions. (a) Schematic illustrating the particle interactions, wall interactions and collision behaviors for the computational model. (b) Schematic of the collective cell migration simulation with confined walls and 600 particles to achieve high-density conditions. Joint distribution of reorientation angle (Δ*θ*_*reorient*_) and inter-cellular deviation angle (*θ*_*between*_) for inelastic collision pairs (c) and elastic collision pairs (d). (e) Representative simulation snapshots, speed and angular velocity at the final time point (*step* = 2000) illustrate how the intermediate fractions of inelastic particles (*p*_in_) exhibit faster and more coordinated motion. (f) Phase diagram of normalized speed and correlation length with varying fraction of particles that exhibit inelastic collisions. Elliptical contours illustrate the Gaussian summaries of the data distribution, constructed from the mean and covariance matrix for *p*_in_ = 0.0-0.2, 0.4-0.6, and 0.8-1.0.

We observed that a homogeneous population of “epithelial” particles exhibited relatively slow migration with larger correlation lengths (Fig. 6ef, Fig. S8b). For a heterogeneous population of equal parts “epithelial” and “mesenchymal” particles, there was faster migration with larger correlation lengths. Finally, homogeneous populations of “mesenchymal” particles exhibited faster migration but with relatively small correlation lengths. Overall, these computational results suggest that “epithelial” cells exhibit collision guidance mechanisms that promote collective migration, but these inelastic collisions gradually slow cells down so they will eventually arrest at high density, consistent with experimental observations. Instead “mesenchymal” cells favored “uninhibited” or “repulsive” elastic interactions, particularly when colliding at large angles. Together, these experimental and computational results demonstrate that EMT state-dependent interactions mediate collective migration. We identified a “collision guidance” behavior that can promote persistent motion with long-range velocity correlations, while “repulsive” or “uninhibited” behavior resulted in more individual migration with diminished coordination.

### E. Partial EMT Monolayer Fronts Overwhelm Control or EMT Monolayers

EMT often occurs at collective invasion fronts [40], and we sought to determine whether difference in cell-cell interactions would have relevance at much larger tissue length scales, such as collisions between expanding tissue monolayers. We prepared tissue monolayers with different EMT states and examined how their respective fronts collided over time. Briefly, we used a poly(dimethylsiloxane) (PDMS) stencil to pattern cells into separated regions. Removal of the stencil allowed the tissue monolayers to expand outward, ultimately resulting in a collision between the two collectively migrating fronts (Fig. 7a). Since these cells were typically near confluency at the start of the assay, migration was primarily driven by expansion of the monolayer, resulting in no significant differences in migration speed 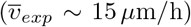 across the TGF-*β*1 conditions (Fig. 7b). In contrast, monolayers prepared using control and 1 ng/mL TGF-*β* treated cells were more coordinated with larger correlation lengths of ∼ 200 *µm*, while monolayers from 5 ng/mL TGF-*β*1 treated cells were less coherent (∼ 130 *µm*) (Fig. 7c). The collision of two untreated monolayers with epithelial cells (Fig. 7d) resulted in immediate arrest of both collective fronts, forming a stable boundary (Fig. 7e) and slowdown of cells on both sides (Fig. 7f). In comparison, the collision between a 1 ng/mL TGF-*β*1–treated monolayer and a control monolayer also resulted in the formation of a stable boundary (Fig. 7g). Unexpectedly, the control cells transiently moved rearward after collision, so that the velocity of the 1 ng/mL TGF-*β*1 treated monolayer propagated across the boundary into the control monolayer (∼ 20 hours) (Fig. 7h). Indeed, we observed periodic, long-range velocity “pulses” with positive peaks observed ∼150*µ*m distant from the collision boundary (Fig. 7h). Similar “deformation wave” dynamics have been reported elsewhere after the collision of epithelial monolayers [37] (Fig. 7k). Finally, the collective front of a 1 ng/mL TGF-*β*1 treated monolayer continued to advance after contacting the 5 ng/mL TGF-*β*1 treated monolayer (Fig. 7ij). The expansion velocity again extended past the boundary, suggesting that the cells of the 5 ng/mL TGF-*β*1 treated monolayer were repelled and migrated backwards. Immunostaining revealed that control cells exhibited high E-cadherin, whereas 1 ng/mL and 5 ng/mL TGF-*β*1 exhibited decreased E-cadherin and increased N-cadherin (Fig. S9). Thus, the 5 ng/mL TGF-*β*1 treated cells likely exhibited weak cohesion and were unable to withstand collisions with another front. Overall, these results show that 1 ng/mL TGF-*β*1 drives cohesive and invasive monolayer fronts that can overcome monolayer fronts from untreated controls or 5 ng/mL TGF-*β*1, consistent with the persistent collective migration behaviors we observed for cells exhibiting partial EMT.

**FIG. 7.**
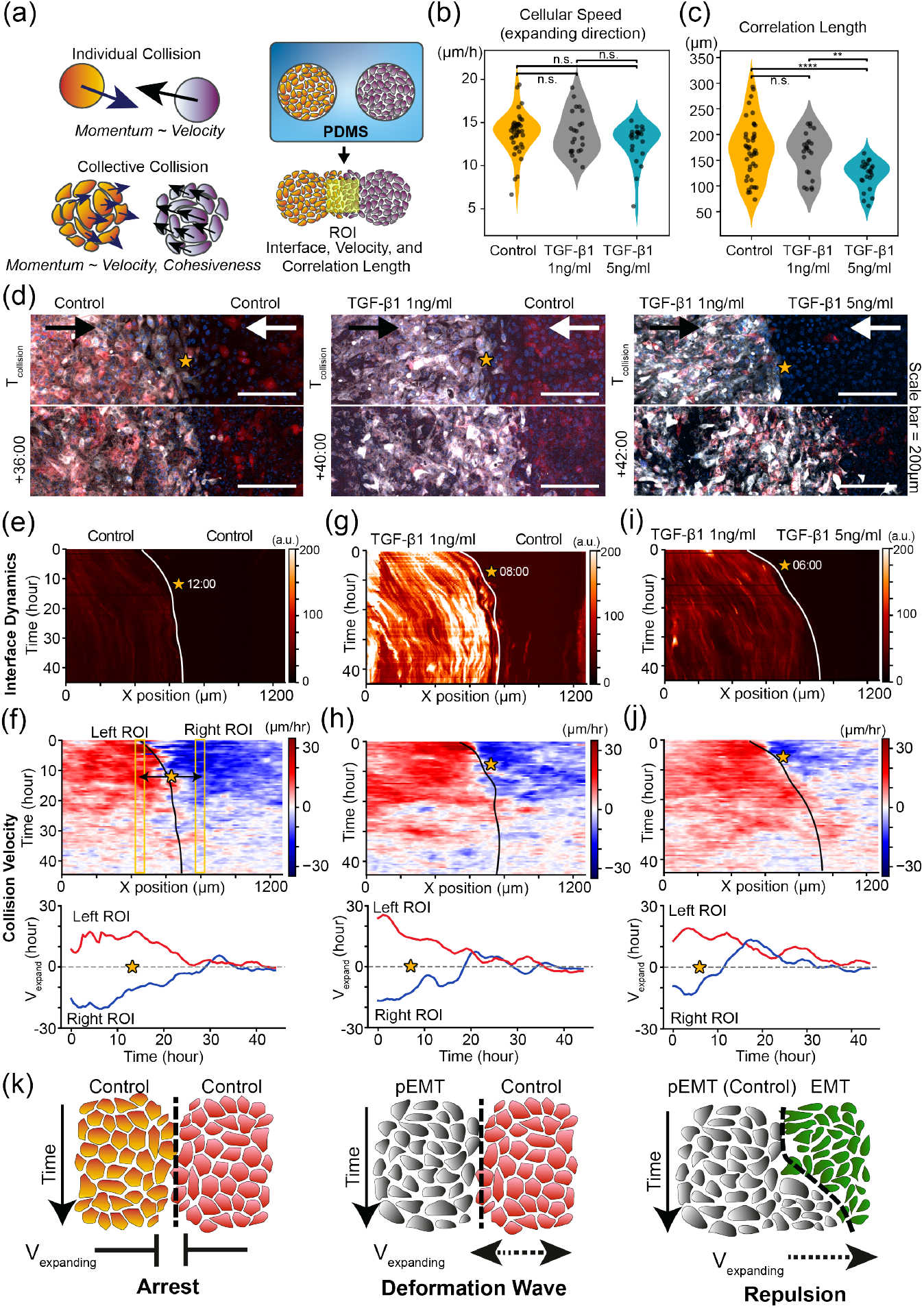
Partial EMT collective fronts drive deformation waves and repulsion after colliding with other cell types. (a) Schematic illustration of collective collision dynamics and experimental setup using a PDMS stencil to induce controlled collective collisions. (b, c) Mean migration speed of expanding direction and correlation length of expanding monolayer across TGF-*β*1 pretreatment conditions (*n* = 3 independent experiments). Statistical comparisons were performed using one-way ANOVA, followed by Welch’s unequal-variance *t*-tests with Bonferroni correction for multiple comparisons. Significance levels are indicated as ^*^*p <* 0.05, ^**^*p <* 0.01, ^***^*p <* 0.001, and ^****^*p <* 0.0001. (d) Representative snapshots of monolayer expansion and collisions between distinct epithelial or EMT states. Kymograph of collective front and cell velocity for control-control collisions (e,f), TGF-*β*1 (1 ng mL^−1^)–control collision (g,h), and TGF-*β*1 (1 ng mL^−1^)–TGF-*β*1 (5 ng mL^−1^) collision (i,j). Orange stars indicate the spatiotemporal position of collective collisions (k) Schematic of collision behaviors associated with arrest, deformation wave, and repulsion.

## III. DISCUSSION

Confluent epithelial monolayers tend to be arrested while dispersed mesenchymal cells exhibit uncoordinated motility [3]. Epithelial cells transition to collective migration during embryonic development, wound healing and tumor invasion [22], which may occur through a partial EMT [1]. In this article, we show that mammary epithelial cells exhibit coordinated and persistent motility after treatment with moderate concentrations of TGF-*β*1. Our single cell measurements using a dual-color fluorescent reporter reveal an increase in population heterogeneity with a shift towards more cells with no fluorescent protein expression (E-cad- / ZEB1-) or green fluorescent protein expression (ZEB1+). Nevertheless, we see the largest alteration in migratory behavior in cells with red fluorescent protein expression (E-cad+), showing increased persistence at high cell densities. One limitation of fluorescent reporters is that there is a time lag for measurable changes in fluorescent protein expression, which may explain the similarities in migration behavior between red (E) and uncolored (P) cells. We have previously observed that EMT induction using TGF-*β* results in heterogeneous kinetics at the single cell level [15], which is consistent with the heterogeneity we observe here in fluorescent reporter expression. In comparison, inducible gene expression systems that promote EMT transcription factors (e.g. Snail) may rapidly and more uniformly drive cells towards a fully mesenchymal state [15]. Thus, it remains challenging to generate a more homogeneous population of cells with partial EMT states, given our limited control over the underlying signaling networks.

We show that pairs of epithelial and partial EMT cells (based on fluorescent reporter expression) exhibit “collision guidance” that facilitates coordinated migration, whereby transiently interacting cells will steer their migration direction along parallel trajectories. Instead, mesenchymal cells are more uninhibited and migrate over each other without changing direction. Moreover, epithelial and mesenchymal cells are more likely to repel upon contact and migrate away from each other (e.g. contact inhibition of locomotion). Notably, partial EMT cells can sustain collision guidance with both epithelial and mesenchymal cells, even for nearly head-on collisions at large angles. These differences in cell-cell response may reflect differences in cell surface receptor expression (e.g. N-cadherin [41], Eph / Ephrins [42], Slit-Robo [43] etc.) (Fig. S9) as well as cytoskeletal regulation of protrusions and lamellipodia. We have developed a minimal physical model based on self-propelled particles that recapitulates these cell-cell interactions via elastic or inelastic collisions. Although this model is sufficient to achieve qualitative agreement with our experimental results, there we have not accounted for certain aspects of phenotypic heterogeneity that would increase model complexity. For instance, we do not consider differences in speed between EMT states, transitions between EMT states, or proliferation. We also limited our model to two cell types, but additional cell types with heterogeneity in surface receptor expression might help to map out the landscape of functional EMT states.

Remarkably, these heterotypic interactions at the single cell level can drive complex larger-scale collision dynamics between expanding monolayer fronts. We find that collective monolayer fronts from cells treated with 1 ng/mL TGF-*β* are able to overcome collective monolayers from untreated cells or cells treated with 5 ng/mL TGF-*β*, propagating long ranged velocity pulses deep into the other monolayer have occurred. Indeed, the untreated epithelial monolayers immediately arrest their migration after collision (consistent with epithelial contact inhibition of locomotion), while the 5 ng/mL TGF-*β* treated monolayers lack cohesion and cannot repel invasion by the 1 ng/mL TGF-*β* treated monolayer. Future work could consider how partial EMT mediates monolayer collision in more complex geometries [39], as well as collective dynamics in spatial confinement [44–46] or 3D matrix [47–50].

In conclusion, we find that mammary epithelial cells exhibit persistent collective migration after treatment with moderate TGF-*β* concentrations, which differs from arrested motility of untreated cells and uncoordinated motility after treatment with high TGF-*β* concentrations. Our single cell analysis with a dual fluorescent EMT reporter shows that treatment with moderate TGF-*β* concentration results in highly heterogeneous populations that include epithelial, partial EMT and mesenchymal states. Nevertheless, epithelial and partial EMT states exhibit cell-cell interactions that promote collective migration by aligning their direction of motion, whereas mesenchymal cells more uninhibited and repulsive interactions. We recapitulate these experimental observations using a minimal physical model of self-propelled particles that undergo either inelastic (epithelial-like) or elastic (mesenchymal-like) collisions. Finally, we show that these differences in cell-cell interactions also result in dramatic differences in tissue-scale interactions, so that collective fronts of partial EMT populations can overcome epithelial or mesenchymal populations. Overall, we show that partial EMT cells interact cooperatively with epithelial or mesenchymal cells to sustain coordinated motility, which is not observed with more homogeneous epithelial or mesenchymal populations. This collective migration is relatively robust to heterogeneity in cell-cell adhesions, which may have implications for understanding how different cell types coordinate their motion when sculpting epithelial tissues and tumors.

## IV. EXPERIMENTAL SECTION

### A. Epithelial Cell Culture

Human mammary epithelial cells (MCF-10A) stably transduced with a Z-CAD dual fluorescent reporter expressing d2GFP and dsRed were a generous gift from M.J. Toneff and J. M. Rosen. The Z-CAD dual sensor consists of a destabilized GFP containing ZEB1 3’ UTR and RFP on the E-cadherin (CDH1) promoter [16]. The MCF-10A cell line was cultured in growth medium made with DMEM / F12 (Invitrogen, 11965-118) with horse serum 5% (Invitrogen, 16050-122), 20ng/ml human epidermal growth factor (R&D Systems, 236-EG), 10 *µ*g/mL insulin (Sigma Aldrich, I-1882), 0.5 *µ*g/mL hydrocortisone (Sigma Aldrich, H-0888), 100 ng/mL cholera toxin (Sigma Aldrich, C-8052), and 1% penicillin streptomycin (Invitrogen No. 15070-063).

EMT was induced in MCF-10A cells using 1 ng/mL or 5 ng/mL TGF-*β*1 (R&D Systems, 240-B002) in growth media. Cells were cultured for 7 days in T25 tissue culture flasks (Genesee Scientific). To maintain cell-cell contact under normal conditions, cells were seeded at a density of 5 × 10^5^ per flask (∼ 25% confluency). For EMT induction with TGF-*β*1, cells were seeded at a lower density of 2 × 10^5^ cells per flask (∼ 10% confluency). This lower initial cell density was intended to keep cells more dispersed and limit the formation of cell-cell junctions that sustain epithelial polarity.

### B. Time-Lapse Live Cell Microscopy

24 h prior to imaging, MCF-10A cells were washed twice with 3.0 mL calcium- and magnesium-free PBS (Cytiva, SH30256), then detached using 1.0 mL Accumax (Sigma-Aldrich, A7089) and 1.0 mL of 0.05% trypsin (Cytiva, SH30042). MCF-10A cells were fluorescently labeled with 1 *µ* M CellTracker^™^ Deep Red (Invitrogen, C34565) in serum-free DMEM/F12 (Invitrogen, 11965-118) to complement fluorescent reporter expression. Cells were incubated for 40 minutes with intermittent gentle mixing by pipetting at 5-minute intervals. Cells were then plated at a density of 2 × 10^4^ cells per well (approximately 60% confluence) in high-content 96-well plates (Corning, 4680) coated with 5 *µ*g/cm^2^ fibronectin (R&D Systems, 1918-FN) to enhance cell adhesion. After seeding, cells were allowed to adhere for 24 h in a humidified incubator at 37°C with 5% CO_2_. Immediately prior to live-cell imaging, nuclei were labeled with 5 *µ*g/mL Hoechst 33342 (Invitrogen, H3570) in PBS for 15 minutes, followed by three washes with serum-free DMEM/F12. Finally, 500 *µ*L of growth medium was added to each well before imaging.

Live cell time-lapse fluorescence microscopy was performed over 24 h using a Nikon Eclipse Ti fluorescence microscope equipped with a motorized stage, direct illumination LED (Thorburn), a multi-channel light source (Lumencor Spectra-X), spinning disk confocal unit (CrestOptics X-Light V2), cMOS camera (Andor Neo), and a 10× Plan Apo objective (NA 0.75). Cells were maintained in a microscope-mounted incubator (In Vivo Scientific) under controlled conditions of 37 °C, 5% CO_2_, and humidification. NIS Element software was used for automated image acquisition. The Hoeschst, GFP, RFP, and Deep Red CellTracker™ were excited and detected at 395/450 nm, 470/535 nm, 555/583 nm and 640/670 nm, respectively. Z-stack images were acquired every 15 or 20 minutes, capturing at least 3 slices with a z-step size ranging from 2.5-5 *µ*m. Each channel of the z-stacks was processed using maximum intensity projection in ImageJ.

### C. Cell Boundary Segmentation

Cell segmentation was conducted using the CellPose algorithm [51, 52], a deep learning-based open-source tool designed to generalize across various cell types and experimental setups. We firstly employed the pre-trained model Cyto3, the super generalist model, available within the CellPose software package (version 3.0.7) to track the cytoplasm channel of cell images. Before loading images into the software, cell images containing separate cytoplasmic and nuclear channels were merged into an RGB format using ImageJ software. The cytoplasmic channel was assigned to the red channel, while the nucleus channel was allocated to the blue channel of the RGB image. For image segmentation, two fluorescence channels were used. The red channel served as the primary segmentation channel, while the blue channel was optionally incorporated to improve boundary detection. A target cell diameter of 40 px (50.8 *µ*m) was specified during segmentation. Following automated segmentation, results were manually reviewed and corrected to resolve errors such as under-segmentation or incorrectly merged cells, ensuring accurate extraction of morphological features for downstream analysis. The segmentation model was subsequently retrained using 97 manually corrected images randomly selected from different time points (0 h, 9 h, and 18 h after imaging initiation).

## D. Cell Shape Feature Extraction and EMT State Classification from Reporter Expression

The shape feature extraction were calculated from the segmented cell mask from previously published methods [53]. In addition to the geometrical analysis, we also utilized the Zernike moments to get rotational invariant features of cell boundary. To estimate the EMT state at the single-cell level, we classified cells according to the ratio of fluorescence intensities within each cell body. As shown in Fig.S2 b, cells displayed heterogeneous color-expression patterns across a broad range of signal ratios. For each fluorescence channel, cell-body intensities were background-corrected using the signal measured outside the corresponding cell mask. After Min–Max normalization, cells with normalized values below 0.05 were defined as non-colored cells. Among the remaining cells, the dominant color was assigned when one channel exhibited an intensity at least twofold greater than the other.

## E. Cell Migration Tracking

Time-lapse image sequences were analyzed to estimate cellular motion fields using the Farnebäck optical flow algorithm applied to the cytoplasmic intensity channel. The local window size for optical flow was optimized (8.89 *µ*m × 8.89 *µ*m) to be sufficiently small to capture cell-scale motions, while remaining robust to large displacements through the use of a coarse-to-fine pyramid approach. This procedure yields a two-dimensional displacement field at each pixel,

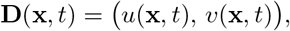

where *u* and *v* denote the horizontal and vertical displacements between consecutive frames. Single-cell trajectories were reconstructed by incrementally updating cell centroid positions using the local displacement field. Cell centroids, **P**_cell_(0), were extracted from segmented cell masks at the initial time point. The centroid position at time *t* was then updated according to

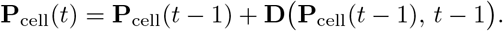

After trajectory reconstruction, spurious trajectories arising from limitations of the optical flow estimation were removed. Trajectories were excluded when frame-to-frame displacements exceeded the reliable tracking range of the algorithm, resulting in loss of correspondence between cell centroids and the displacement field, or when trajectories became trapped in background regions or persisted near image boundaries due to tracking failure. Figure S2 demonstrates that the optical flow-based algorithm produced tracking errors at the length of subcellular level when compared to manually tracked cell trajectories.

### F. Extracting Quantitative Metrics from Cell Trajectories

We used various quantitative metrics to analyze the single cell trajectories in 8 h intervals. Briefly, cell trajectories were represented as centroid coordinates **r**_*t*_ = (*x*_*t*_, *y*_*t*_) at each time frame(*t*). Each trajectory was independently analyzed to extract its kinematic, geometric and coordination-related motility features.

The instantaneous displacement between successive frame was defined as, *d*_*t*_ = ∥**r**_*t*_ − **r**_*t*−1_ ∥, and the instantaneous speed was computed as *v*_*t*_ = *d*_*t*_/Δ*t*, where Δ*t* denotes temporal interval. The cumulative path length was calculated as, *L* = ∑_*t*_ *d*_*t*_, while the euclidean distance from the initial position was defined as, *D*_*t*_ =∥ **r**_*t*_ − **r**_0_∥ . From these two path-related quantities, we derived the directionality *M* = *D*_*t*_/*L* and the outreach ratio *O* = max(*d*_*t*_)/*L*, which characterize directional persistence and step-wise dominance, respectively. Arrest behavior was quantified using an arrest coefficient, defined as the fraction of trajectory steps with *d*_*t*_ below a predefined displacement threshold (5 pixels). The geometry of trajectories was quantified using a loop score defined as the Loop-like trajectory geometry was further quantified using a loop score defined as the ratio between net displacement and total path length(directionality for total time frame), 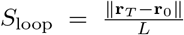. The spatial coverage of each trajectory was quantified by the polygon area enclosed by the ordered centroid coordinates, computed using the shoelace formula.

Directional fluctuations were characterized by turning angles computed from successive displacement vectors. For each interior trajectory point, the turning angle was defined as Δ*θ*_*t*_ = *θ*_*t*+1_ − *θ*_*t*_, when *θ*_*t*_ = atan2(*y*_*t*_ − *y*_*t*−1_, *x*_*t*_ − *x*_*t*−1_). From the distribution of Δ*θ*_*t*_L, we calculated the angular variation (standard deviation of Δ*θ*_*t*_), a turning index defined as, *T*_*i*_ = |∑_*t*_ Δ*θ*_*t*_| /(2*π*), and a curvature count corresponding to the number of turning events with|Δ*θ*_*t*_| *> π/*2.

To assess coordination with neighboring cells, we quantified a velocity correlation metric based on cosine similarity of displacement vectors. For each evaluated frame, neighboring cells within a radius *R* = 100 pixels (127*µm*) were identified, and up to *k* nearest neighbors were selected. The cosine similarity between the displacement vector of the reference cell and that of each neighbor was computed as

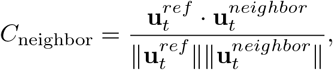

where **u**_*t*_ = **r**_*t*_ − **r**_*t*−1_. Velocity correlation was averaged across neighbors and time points. To reduce computational cost while preserving temporal trends, this calculation was performed every *s* frames (typically *s* = 5), and frames lacking valid neighbors or neighbor history were excluded from the averaging. Together, this framework provides a comprehensive quantification of single-cell motility, capturing instantaneous kinematics, long-range path geometry, directional variability, arrest dynamics, and local coordination within migrating cell populations.

### G. UMAP embedding of single-cell motility features

To characterize the heterogeneity of single cell migration behaviors, the 17 migration metrics for each 8 h interval was represented as a feature vector and standardized by z-score normalization across the full dataset. Dimensionality reduction was performed using Uniform Manifold Approximation and Projection (UMAP) to embed the high-dimensional motility feature space into two dimensions, preserving local neighborhood relationships between trajectories. UMAP was applied with a fixed random seed to ensure reproducibility, and the number of nearest neighbors and minimum distance parameters were selected to balance local structure preservation and global continuity (number of neighbors = 30, minimum distance = 0.3). The resulting low-dimensional embedding was used as a descriptive manifold to visualize the organization of motility phenotypes across experimental conditions and time segments. To quantify temporal evolution, population centroids were computed for each condition and time segment, and centroid displacement vectors were calculated between adjacent segments. Similarity between distributions was assessed using pairwise overlap analysis on the embedded space. For each pair of groups, probability density functions were estimated using Gaussian kernel density estimation (KDE) on a shared grid of data. The densities were normalized to unit integral, and the overlap coefficient (OVL) was computed as

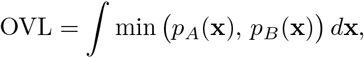

which was numerically approximated using Riemann summation over the discretized grid. Comparisons with insufficient sample sizes were excluded.

**TABLE I.** Summary of single-cell motility metrics extracted from trajectory analysis.

| Category | Metric | Definition |
| --- | --- | --- |
| **Kinematic** | Instantaneous displacement ($d_t$) | $d_t = \ \mathbf{r}_t - \mathbf{r}_{t-1}\ $ |
| | Instantaneous speed ($v_t$) | $v_t = d_t / \Delta t$ |
| | Cumulative path length ($L$) | $L = \sum_t d_t$ |
| | Euclidean distance ($D_t$) | $D_t = \ \mathbf{r}_t - \mathbf{r}_0\ $ |
| | Maximum step size | $\max(d_t)$ |
| | Mean squared step (MSD) | $\frac{1}{N} \sum_t d_t^2$ |
| | Average acceleration | $\left\langle \left\vert \frac{\Delta v_t}{\Delta t} \right\vert \right\rangle$ |
| **Path geometry** | Directionality ($M$) | $M = D_T / L$ |
| | Outreach ratio ($O$) | $O = \max(d_t) / L$ |
| | Loop score ($S_{\text{loop}}$) | $S_{\text{loop}} = \frac{\ \mathbf{r}_T - \mathbf{r}_0\ }{L}$ |
|  | Spatial coverage | Polygon area enclosed by ordered centroid coordinates (shoelace formula) |
| **Directional variability** | Turning angle ($\Delta\theta_t$) | $\Delta\theta_t = \theta_{t+1} - \theta_t, \quad \theta_t = \text{atan2}(y_t - y_{t-1}, x_t - x_{t-1})$ |
| | Angular variation | $\text{std}(\Delta\theta_t)$ |
| | Turning index ($T_i$) | $T_i = \vert \sum_t \Delta\theta_t\vert / (2\pi)$ |
| | Curvature count | Number of events with $\vert \Delta\theta_t\vert > \pi/2$ |
| | Turn count | Number of heading changes with $\vert \Delta\theta_t\vert > 60^\circ$ |
| **Arrest dynamics** | Arrest coefficient | Fraction of steps with $d_t$ below predefined threshold |
| **Coordination** | Velocity correlation ($C_{\text{neighbor}}$) | $\frac{\mathbf{u}_t^{\text{ref}} \cdot \mathbf{u}_t^{\text{neighbor}}}{\ \mathbf{u}_t^{\text{ref}}\ \ \mathbf{u}_t^{\text{neighbor}}\ }, \quad \mathbf{u}_t = \mathbf{r}_t - \mathbf{r}_{t-1}$ |

### H. Optical-flow velocity fields and correlation length analysis

Cell motion fields were estimated from time-lapse image sequences using dense optical flow. For each pair of consecutive frames, optical flow was computed with the Farnebäck algorithm on the red intensity channel, yielding a two-dimensional displacement field **v**(**x**) = (*v*_*x*_(**x**), *v*_*y*_(**x**)) at each pixel **x**. The local speed magnitude was defined as 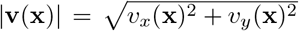. For reporting an average motility scale per frame, mean speed was computed over “active” pixels satisfying **v**(**x**) *v*_min_ ≥ (*v*_min_=1.0 pixel/frames, 5.08*µ*m/hour).

To quantify the spatial coordination of motion, we computed a vector-field correlation length using a Fourier-based autocorrelation approach. For a given region of interest (ROI), the velocity autocorrelation field was obtained as *C*(Δ**x**) = ℱ^−1^ (|ℱ (*v*_*x*_)|^2^) + ℱ^−1^ (|ℱ (*v*_*y*_|^2^), where ℱ and ℱ^−1^ denote the two-dimensional Fourier transform and its inverse, respectively, and the correlation field was centered using an FFT shift. The correlation map was normalized by its maximum value, *C*(Δ**x**) ← *C*(Δ**x**)/ max_Δ**x**_ *C*(Δ**x**), and converted to an isotropic radial profile by averaging over all pixels at the same radius, yielding *C*(*r*) as a function of radial distance *r*. Correlation length was defined as the characteristic decay scale at which the normalized radial correlation dropped below 1/*e*, i.e., *ξ* was the smallest *r* satisfying *C*(*r*) ≤ 1/*e*. Distances were converted to physical units using a pixel-to-length factor (e.g., 1.27 *µ*m per pixel). To reduce bias from spatial inhomogeneity and background regions, the full-frame velocity field was analyzed using a sliding-window scheme. The image was tiled into square sub-windows (e.g., 256 × 256 pixels) with a fixed stride (e.g., 64 pixels). For each window, an “active fraction” was computed as the fraction of pixels with |**v**(**x**) |*v*_min_ (*v*_min_=1.0 pixel/frames, 5.08*µ*m/hour). Windows with an active fraction below the threshold (80%) were excluded to calculate the correlation length only for confluent layer, and the frame-level correlation length was reported as the mean of the correlation lengths computed across all retained windows.

### I. Patch-based quantification of local cellular heterogeneity

Local phenotypic heterogeneity (in color expression and morphology) was quantified using a patch-based analysis. For each sample and time point, cell centroids (*x*_*i*_, *y*_*i*_) were partitioned into non-overlapping square patches of side length *S* = 75 pixels (95.3 *µ*m). Then, patches containing fewer than six cells were excluded. Categorical states were embedded into fixed two-dimensional coordinates. Color identities were mapped as Green → (1, 1), Red → ( −1, −1), Yellow → (−1, −1), and Black → (0, 0). Morphological shape clusters (0–2) were placed at the vertices of an equilateral triangle:

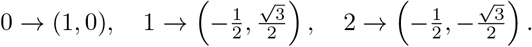

Within each patch containing *n* cells, heterogeneity was defined as the mean pairwise Euclidean distance among embedded coordinates:

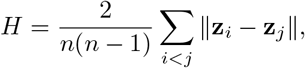

where **z**_*i*_ denotes either the color or cluster embedding. Color and cluster heterogeneity were computed separately and summed to obtain a combined heterogeneity score. Patch-level motility metrics were calculated as the mean and median across cells within each patch.

### J. Quantification of cell–cell interaction behaviors

Cell–cell collision events were identified by integrating cell-resolved trajectories with segmentation masks. First, potentially interacting cell pairs were identified based on spatial proximity. For each frame, all cells present in the field of view were queried for neighbors within a proximity radius of 50 pixels, and unique cell pairs were retained. For each pair, the intercellular distance *d*(*t*) was computed over the time interval during which both trajectories coexisted, and the collision time *t*_*c*_ was defined as the frame at which *d*(*t*) reached its minimum value.

To focus on transient collision events rather than persistent neighboring behavior, cell pairs that remained within the proximity radius for extended durations (*>* 24 frames) were excluded. In addition, candidate events were filtered based on pre-collision motility. For each cell, a pre-collision displacement vector **v**^pre^ was computed over a temporal window of *dT* = 6 frames ending at *t*_*c*_. Candidate pairs were discarded when both cells exhibited negligible pre-collision motion, defined as a displacement magnitude smaller than 2.5 pixels over the pre-collision window.

Physical contact between cells was confirmed using segmentation masks at the collision time *t*_*c*_. Binary masks corresponding to each cell were converted into boundary maps using a morphological boundary detection algorithm and subsequently dilated by one pixel to account for segmentation uncertainty. A collision event was retained only when the overlap between the dilated boundary maps of the two cells exceeded a predefined threshold (three pixels), ensuring direct cell–cell contact.

Collision-induced cellular responses were quantified by comparing pre- and post-collision displacement vectors. For each cell, pre- and post-collision displacement vectors, **v**^pre^ and **v**^post^, were computed over symmetric windows of *dT* = 6 frames before and after *t*_*c*_, respectively. The reorientation angle of each cell was defined as Δ*θ*_reorient_ = cos^−1^(**v**^pre^ · **v**^post^/(∥**v**^pre^∥ ∥**v**^post^∥)). The relative alignment between the two cells after collision was quantified as the intercellular angle 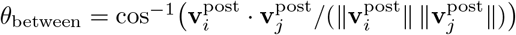.

To distinguish repulsive from following-like responses, an approach axis **ê**_app_ was defined as the unit vector connecting the centroids of the two cells at the collision time *t*_*c*_. The incidence and reflection angles of the focal cell were computed as *θ*_inc_ = cos^−1^(**v**^pre^ · **ê**_app_/ ∥**v**^pre^∥) and *θ*_ref_ = cos^−1^(**v**^post^ **ê**_app_/∥ **v**^post^ ∥), respectively. The absolute angular difference Δ*θ*_diff_ = |*θ*_inc_ − *θ*_ref_ | was used as a quantitative measure of collision-induced angular deviation. Distributions of collision responses were summarized using kernel density estimation and stratified according to EMT reporter state and morphological class.

### K. Simulations of self-propelled particles with elastic or inelastic collisions

We modeled epithelial or mesenchymal cells as self-propelled particles with distinct interaction rules to investigate the emergence of collective migration. Collision interactions were evaluated for all particles pairs within a contact distance *r*_coll_ = 2.0 using a grid-based neighbor search, and overlaps were resolved by a small symmetric position correction to prevent overlap. Type-dependent collision responses were then applied: Epithelial–epithelial collisions were modeled as one-shot perfectly inelastic events in which both particles adopted the center-of-mass velocity when approaching; Mesenchymal–mesenchymal collisions were treated as elastic normal impulses with restitution coefficient *e*_*GG*_ = 1.0; and epithelial–mesenchymal collisions were treated as elastic impulses using the same restitution rule as mesenchymal-mesenchymal collisions. A pairwise cooldown gate was included (set to zero in the reported runs) to prevent repeated collision-triggering within a short interval. In addition, Vicsek alignment was applied with type-specific neighborhood radii (*r*_vicsek,*R*_ = 2.0, *r*_vicsek,*G*_ = 2.0) and alignment strength *α*_*v*_ = 0.10; for each particle, the velocity direction was partially updated toward the average direction of neighboring particles within the corresponding radius, while preserving its speed.

Particle motion was governed by overdamped active dynamics driven by persistent stochastic forcing and confinement. Specifically, velocities were updated from an acceleration term composed of persistent Ornstein–Uhlenbeck (OU) acceleration noise with type-specific parameters (*τ, σ*) (here *τ*_*R*_ = *τ*_*G*_ = 1.0 and *σ*_*R*_ = *σ*_*G*_ = 0.7). Positions were then advanced by explicit Euler integration.

A total of *N* = 600 particles were initialized with uniformly random positions inside a circular domain of radius *R*_domain_ = 30 and random initial velocity directions with magnitude *v*_init_ = 1.0. If particles moved outside this domain, they experienced a soft wall repulsion to weakly reproject them inward, implemented as an inward acceleration proportional to the radial overshoot with strength *k*_wall_ = 50. The fraction of mesenchymal particles *p*_*in*_ was varied from 0 to 1 in increments of 0.1, and each condition was repeated for five independent runs using different random seeds. Simulations were integrated with timestep *dt* = 0.05 for *T* = 2000 steps, while enforcing a maximum speed *v*_max_ = 2.5.

To quantify collective migration, simulations were sampled every 50 steps. The mean speed was computed as the population-average of |**v**|, and global polarization was computed as 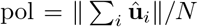, where 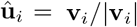. A correlation length was estimated from random pairs of particles by binning the pairwise velocity dot product 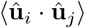 as a function of inter-particle distance *r* and defining *ξ* as the first distance where the binned correlation decayed below 1/*e* of its near-field value. Time series and last-frame summary metrics were recorded for each run and used to construct phase diagrams as a function of *p*_red_.

### L. Tissue Expansion Assay

To conduct collective collisions between epithelial cell layers, we utilized a polydimethylsiloxane (PDMS) stencil to partition different cell conditions and initiate tissue expansion after the cells reached a confluent layer. We used a 10:1 ratio of elastomer and cross-linker (SYLGARD 184, Dow) to fabricate a silicone sheet with a height of 5 mm. After solidifying the silicone sheet for 2 hours at 60 °C, circular holes were punched using a 4 mm biopsy punch (Robbins Instrument) to create wells for the cell layers, and a 10 mm biopsy punch was used to cut the overall stencil structure. Each stencil was sterilized in 70% ethanol for 30 minutes, followed by 1 hour of ultraviolet (UV) exposure in a biosafety cabinet. The stencils were then washed three times with PBS (Cytiva, SH30256) and coated with a 3% bovine serum albumin (BSA) solution in PBS for 1 hour at 37 °C to prevent undesired attachment between the stencil and cells. After coating, the stencils were washed three times with PBS, thoroughly dried, and placed onto a 24-well tissue culture plate (Corning, 353226). Cells were seeded at 2 × 10^4^ cells per well, and one side of the cells was stained with 1 *µ*M CellTracker^™^ Deep Red dye (Invitrogen, C34565) to visualize interface dynamics. After allowing 24 hours for cell attachment, the stencil was carefully removed vertically to avoid undesired damage to the cells, and live-cell imaging was performed for 48 hours.

### M. Immunofluorescence Staining

To qualitatively assess E- and N-cadherin expression in the cell layer, cells were washed with 1X phosphate-buffered saline (PBS, Cytiva, SH30256) containing calcium chloride and magnesium chloride, and then fixed with 4% paraformaldehyde (Fisher Scientific) in PBS for 20 min at room temperature. The cells were then washed three times for 5 min each with PBS containing 1% bovine serum albumin (BSA, Sigma-Aldrich, A9418), followed by overnight incubation at 4°C with primary antibodies against E-cadherin (Mouse hoset species, BD Bioscience, 610181) and N-cadherin (Rabbit host species, Abcam, ab18203) diluted in PBS containing 1% BSA (1:500 and 1:200, respectively). On the following day, cells were washed three times with 1% BSA solution and incubated in the dark for 2 h on a shaker with goat anti-mouse and goat anti-rabbit secondary antibodies (Alexa Fluor 488 and 647, Thermo Fisher, respectively). After three additional washes with PBS, Hoechst (Invitrogen, H3570) was added to stain nuclei, and the cells were washed once more with 1X PBS before imaging. Fluorescence images were acquired using a 12-bit sCMOS camera (Andor Neo), a 20 × Plan Apochromat objective (NA 0.75, 1 mm working distance), and a light-guide-coupled Lumencore Sola white light excitation system. For throughput, an array of images was acquired for each multiwell plate, while imaging parameters were kept constant across all images and experiments.

### N. Statistical Analysis

All statistical analyses were performed independently for each defined time window and experimental condition. Biological replicates were defined as independent live-imaging sequences. Data normality was assessed using the Shapiro-Wilk test before to selecting parametric or non-parametric statistical comparisons. For normally distributed variables, group differences were evaluated using one-way analysis of variance(ANOVA), followed by Tukey’s post hoc test. For non-normally distributed data, the Kruskal-Wallis test was used, followed by Dunn’s post hoc test with Bonferrnoi correction. For morphology-based comparisons among shape classes, one-way ANOVA with Tukey’s post hoc correction was applied (Fig. S4ab). Quintile-stratified analyses (bottom 20% to top 20%) were assessed using Kruskal–Wallis tests with Holm-adjusted pairwise comparisons (Fig. S4fg). For patch-based heterogeneity stratification (bottom 33% vs top 33%), group differences were evaluated using the Mann–Whitney U test(Fig. S5d,e).

## ACKNOWLEDGMENTS

We thank M.J. Toneff and J.M. Rosen (Baylor College of Medicine) for the kind gift of MCF-10A cells transduced with the Z-cad fluorescent reporter. We also thank D. Bhaskar for helpful discussions on image processing. We acknowledge funding from NIH through Grant R01GM140108 (HJ, JK, IYW), NSF through Grant 2428662 (SEL), and Brown University’s Hibbitt Engineering Postdoctoral Fellowship (JK).

## Author Contributions

HJ, JK, JYS, and SEL performed experiments. HJ, JK, SEL and IYW analyzed data. HJ designed and implemented the computational model. HJ and IYW wrote the paper with feedback from all authors. IYW supervised the project.

## Data Availability Statement

Source data, image analysis and simulation code will be made available from the corresponding author upon reasonable request.

