## Supplementary Information for "Partial EMT Drives Persistent Collective Migration via Collision Guidance in Heterogeneous Populations"

Hyuntae Jeong,[1] Jiwon Kim,[1] Jea-Yun Sim,[1] Susan E. Leggett,[2] and Ian Y. Wong[1]
[1] *School of Engineering, Legoretta Cancer Center. Brown University. Providence RI 02912, USA*
[2] *Department of Bioengineering, University of Illinois Urbana-Champaign. Urbana, IL 61801 USA*

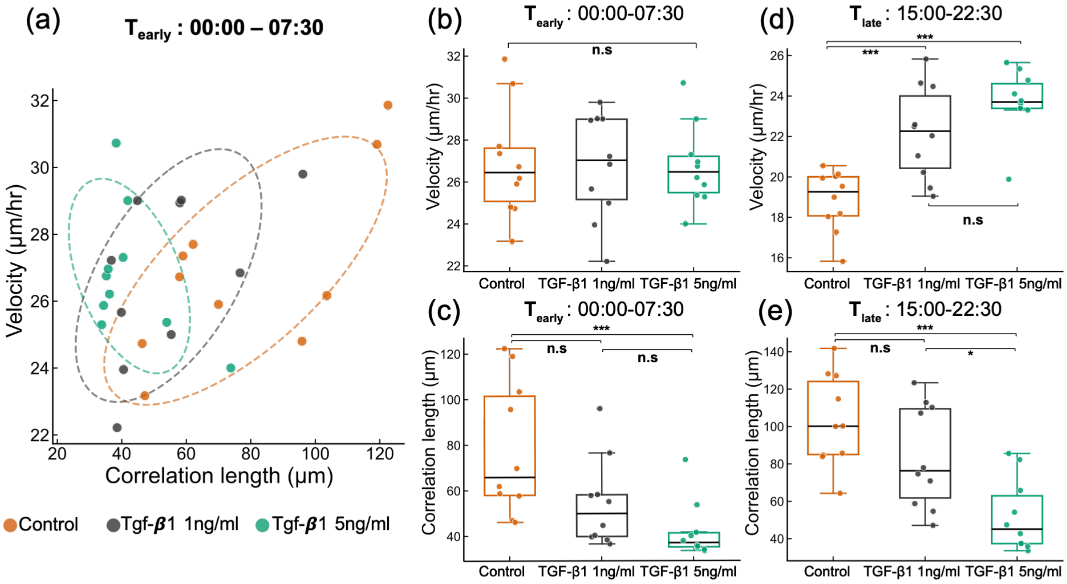


FIG. S1. **Comparison of migration speed and correlation length for untreated controls relative to TGF-$\beta$1 treatment.** (a) Comparison of average correlation length and velocity for untreated controls relative to 1 ng/mL or 5 ng/mL TGF-$\beta$1 treatment. Each scatter point corresponds to one sample within the given time window (0–7.5 h). Elliptical contours illustrate the Gaussian summaries of the data distribution, constructed from the mean and covariance matrix. (b,c,d,e) Box plots show the distributions of cellular speed (b,d) and correlation length (c,e) across conditions during the early(00:00 - 07:30; b,c) and late time(15:00-22:30; d,e) windows. Boxes represent the interquartile range, the center line indicates the median.Individual data points correspond to the one sequence of live image. Statistical comparisons were performed independently for each time window. One-way ANOVA with Tukey's HSD post-hoc test was used when normality assumptions were satisfied(in this case the cellular speed); otherwise, the Kruskal–Wallis test followed by Dunn's post-hoc test with Bonferroni correction was applied(for correlation length).

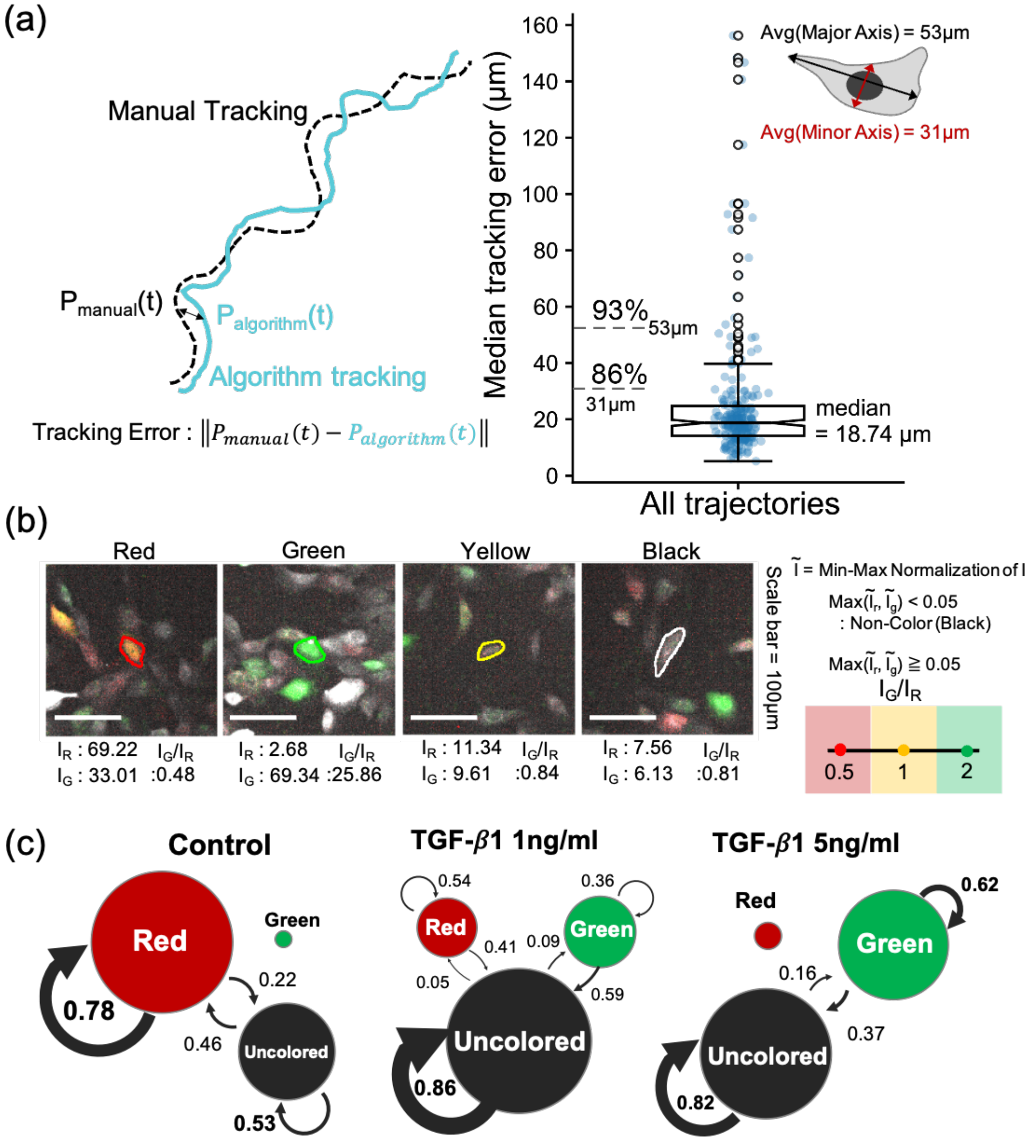


FIG. S2. **Validation of optical flow tracking of cells and annotation method for color reporter expression.** (a) Manually tracked trajectories with trajectories were generated by the algorithm. A total of 246 cells were manually tracked, randomly selected with equal representation from each experimental sample. Tracking error was quantified as the median positional difference between the two trajectories over time. Because manual tracking was performed without using centroids derived from cell masks, an initial positional discrepancy of approximately $\sim 10\ \mu m$ was observed. The median tracking error was 18.74 $\mu$m, which is smaller than the average minor axis length of the cell body (31 $\mu$m) measured from the masks. This indicates that most trajectory discrepancies occurred within the spatial scale of individual cell bodies. The percentages shown in the figure indicate the fraction of trajectories with smaller error than minor axis(81%) and major axis of cells(93%). (b) Representative images for expression of reporter colors and method used to annotate the cell color by measuring RFP to GFP signal ratio. (c) Markov chain of color expression states over 12 hours of cell trajectories. To reduce classification error, a color change was counted only when the new color state was maintained for at least 3 consecutive frames. Circle size is proportional to the fraction of cells in each color state. Arrow width represents the fraction of each transition among all observed transitions, and the number beside each arrow indicates the transition probability.

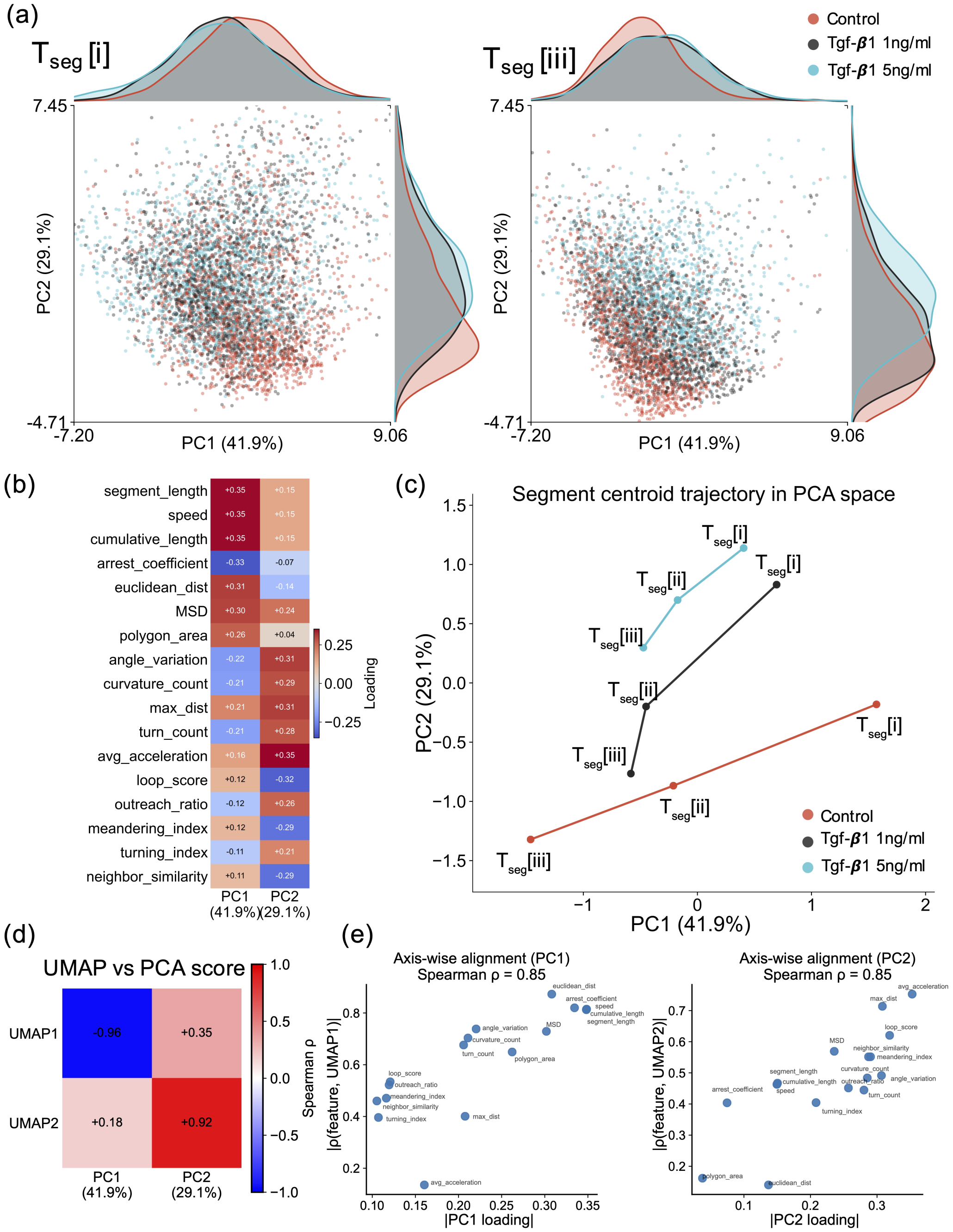


FIG. S3. **Principal component analysis (PCA) shows that single cell trajectories after 1 ng/mL TGF-$\beta$1 treatment are relatively fast and persistent at later times relative to untreated controls or 5 ng/mL TGF-$\beta$1 treatment.** (a) PCA dimensionality reduction of 17 motility metrics for each 7.5 h segment projected into two dimensions. Each point represents an individual cell trajectory, colored by experimental condition (Control, red; TGF-$\beta$1 1 ng ml$^{-1}$, gray; and TGF-$\beta$1 5 ng ml$^{-1}$, cyan). The distributions of trajectories at time segments $T_{seg}$[i] and $T_{seg}$[iii] are shown. Kernel density distributions of each condition along the PC1 and PC2 components are shown in the top and right panels, respectively. (b) Loading of 17 motility features to PC1, PC2 components. PC1 correlates with slow to fast migration, while PC2 correlates with persistent to random migration. (c) Time segment−wise centroid displacement in the PC latent space for each experimental condition. Trajectories indicate the average displacement of population centroids between adjacent segments, revealing directional evolution of motile functions. (d) Spearman rank correlations between UMAP coordinates and PCA scores, (e) Absolute PC1 (left) and PC2 (right) loadings were compared with absolute Spearman correlations of features corresponding to UMAP axes.

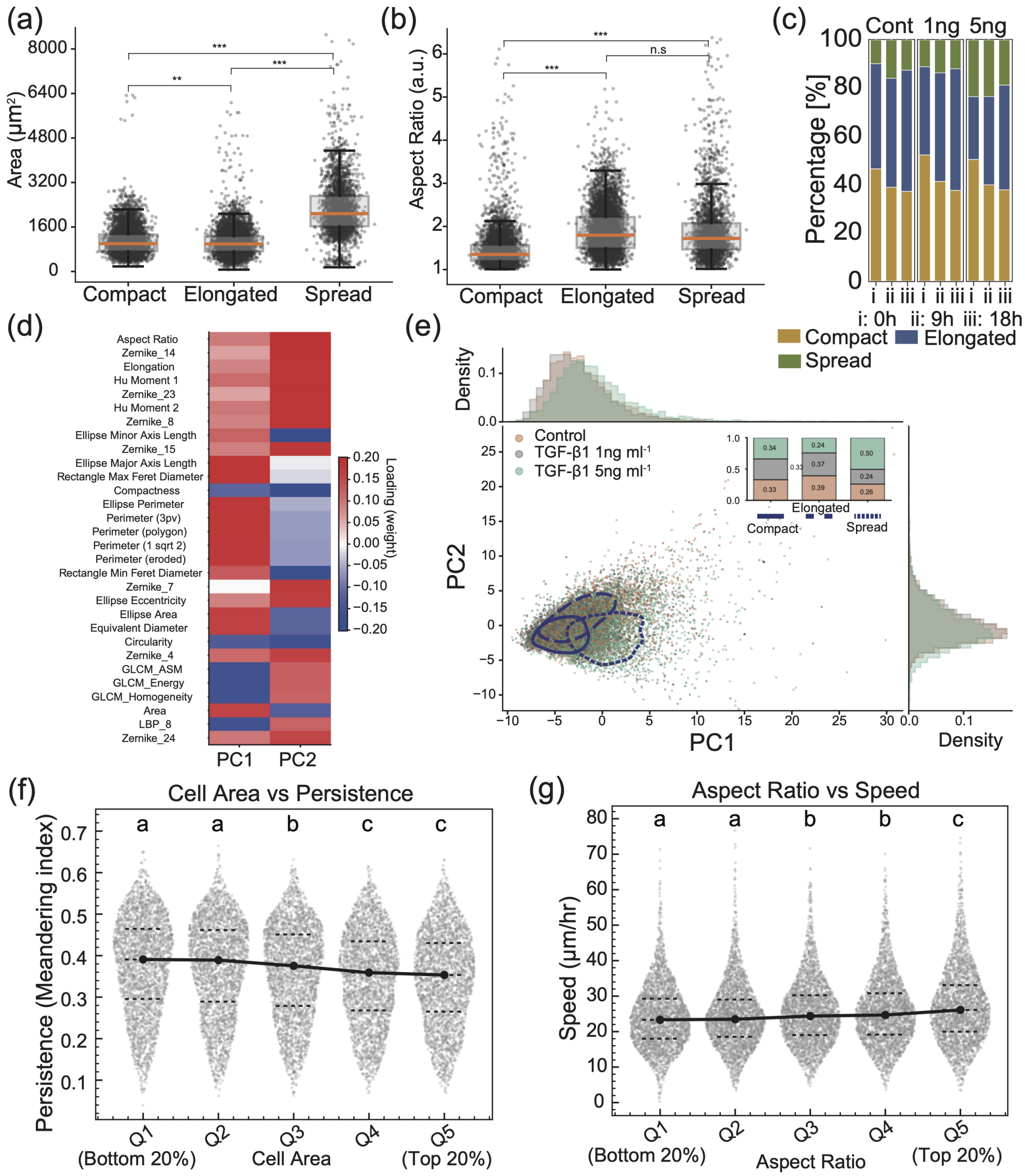


FIG. S4. **Single cell morphology distributions shown in PCA space and relation to motility metrics.** (a,b) Box-and-whisker plots showing the distributions of cell *Area* (a) and *Aspect Ratio* (b) for three types of morphology: Circular, Elongated, and Spreading. Boxes indicate the interquartile range with the median shown as a horizontal line, and whiskers denote 1.5 times the interquartile range.Statistical significance was assessed using one-way ANOVA followed by Tukey's post hoc test; n.s., not significant; $^{**}p < 0.01$; $^{***}p < 0.001$.(c) Percentage of each cell morphology at 0, 9, and 18 hours in untreated control relative to TGF-$\beta$1 concentrations. (d) Heatmap of PCA loadings for the first two principal components (PC1 and PC2), highlighting the top morphology-related features ranked by absolute loading magnitude. Positive and negative loadings indicate features contributing in opposite directions along each principal component, revealing that PC1 is primarily associated with size- and elongation-related descriptors, whereas PC2 captures shape anisotropy and texture-related features. (e) Two-dimensional PCA scatter plot of single-cell morphological features. Each point represents an individual cell and is colored by experimental condition (Control, TGF-$\beta$1 1 ng ml$^{-1}$, and TGF-$\beta$1 5 ng ml$^{-1}$). Smoothed kernel density estimation (KDE)-based contours delineate the high-density regions of each shape class. Marginal histograms along the top and right axes show the condition-dependent distributions of PC1 and PC2, respectively. The inset stacked bar plot summarizes the proportion of shape classes within each experimental condition. (f) Persistence decreases modestly with increasing cell area, whereas (g) migration speed increases with aspect ratio. Cells were grouped into quintiles (bottom 20%–top 20%). Dots represent individual cells, jittered proportionally to local density. Dashed lines indicate quartiles and filled circles represent group means. Statistical significance was assessed using Kruskal–Wallis with Holm correction; groups sharing letters are not significantly different.

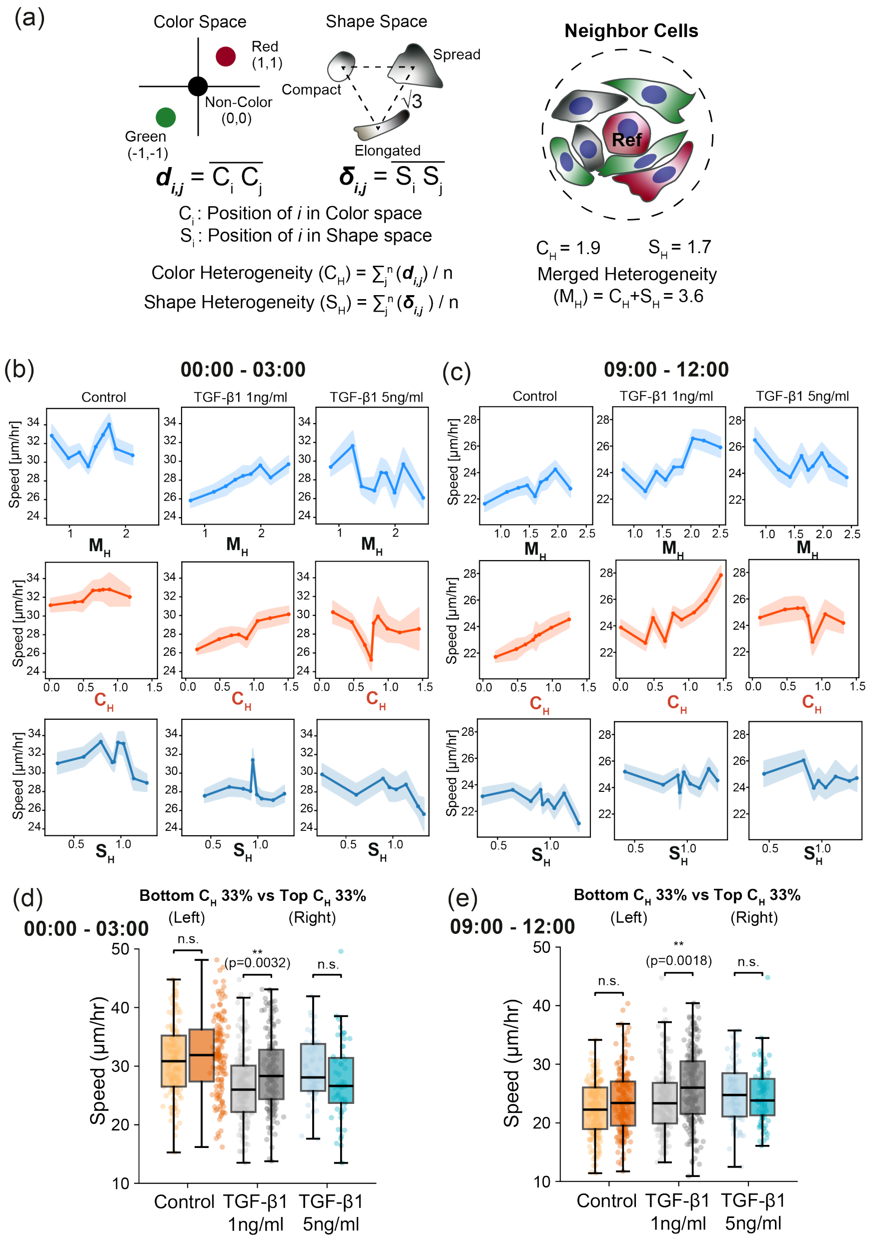


FIG. S5. **Comparison of local heterogeneity based on color expression and shape with average cell speed** (a) Schematic for calculating the local heterogeneity in the patch of cell imaging by projecting the color and shape into a coordinate system established based on their differences. (b, c) Relationship between various local heterogeneity readouts (merged heterogeneity ($M_H$), color heterogeneity ($C_H$), and shape heterogeneity ($S_H$)) and cell speed at early (b) (00:00–03:00) and intermediate (c) (09:00–12:00) across TGF-$\beta_1$ treatment conditions. Data were equally binned by sample count. Markers denote bin-averaged values, and shaded bands represent SEM. (d, e) Patches were stratified into bottom 33% and top 33% of the color heterogeneity index at each time point. Across conditions (Control: orange, TGF-$\beta$1 1ng/ml pretreatment: gray, TGF-$\beta$1 5ng/ml pretreatment: cyan), high-heterogeneity patches display condition-dependent shifts in motility relative to low-heterogeneity patches. Statistical differences were evaluated using permutation testing.

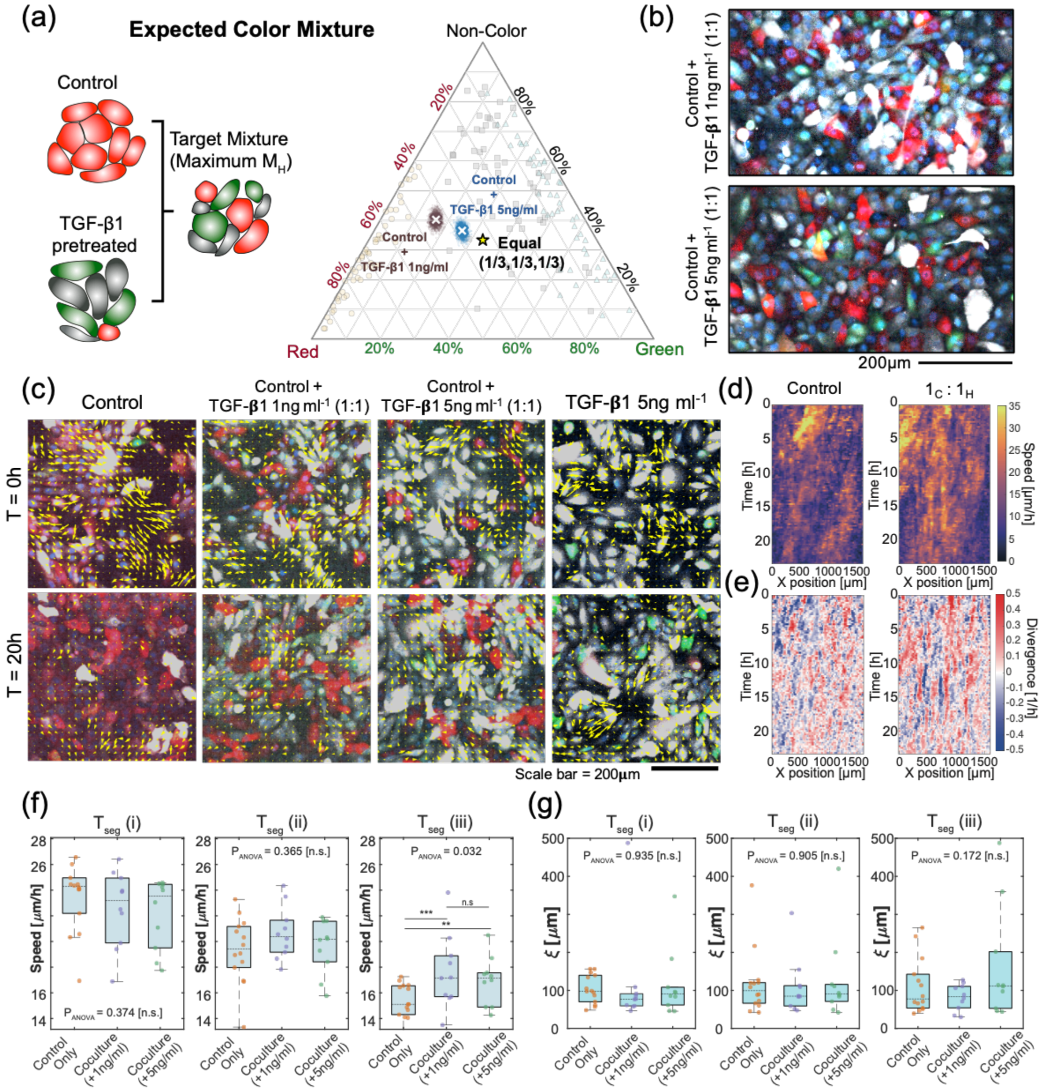


FIG. S6. **Enhanced collective cell migration in co-cultured mixtures of control and TGF-$\beta$1 treated cells.** (a) Approach and expected composition of mixtures from control and TGF-$\beta_1$–treated conditions. (b) Representative snapshots of co-cultured MCF10A monolayers with mixed EMT states (50:50 control and TGF-$\beta_1$–treated cells). Cell cytoplasm is shown in white and nuclei in blue. Red and green indicate reporter-defined cell states. (c) Representative snapshots of migrating cells from co-cultured MCF10A monolayers with mixed EMT states (100:0, 50:50 control and TGF-$\beta$1–treated cells). Images were acquired at 0 and 20 h. Cell cytoplasm is shown in red and white, nuclei in blue, and optical flow–derived migration vectors are shown in green. (d, e) The kymographs for the cellular speeds(d) and divergence(e) of mean value of central 10 grids (1 grid = 16 pixels) over time for control only condition and control + TGF-$\beta$1 5 ng ml$^{-1}$ co-culture condition. (f, g) Quantitative comparison of cellular speed(f) and correlation length(g) in co-cultured conditions. Each dot corresponds to one independent sample, and boxplots show the median and interquartile range (25th–75th percentile).Statistical significance is indicated as $p < 0.05$ (*), $p < 0.01$ (**), and $p < 0.001$ (***).

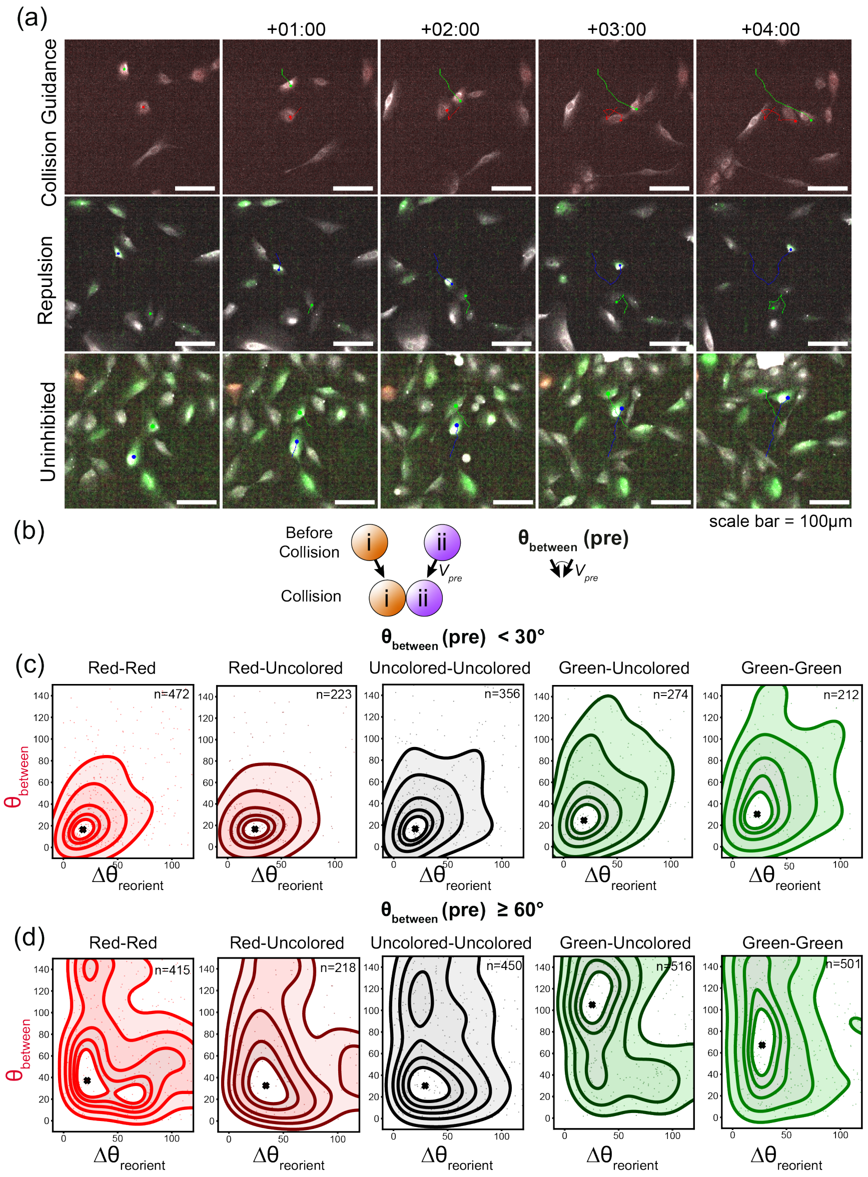


FIG. S7. **Comparison of cell-cell interaction for different fluorescent reporter expression** (a) Representative snapshots for collision guidance, repulsion and uninhibited interactions, (b) Illustration for measuring $\theta^{pre}_{between}$, (c,d) Kernel density estimate contour lines (level = 0.4, 0.6, 0.8, 0.9,and 0.95) of collision-induced angular responses, plotted as the reorientation angle ($\Delta\theta_{\rm reorient}$) versus the intercellular angle ($\theta_{\rm between}$) Each plot shows a subset of data associated with smaller or larger values of $\theta^{pre}_{between}$ ((c):$< 30$ and (d):$\geq 60$)

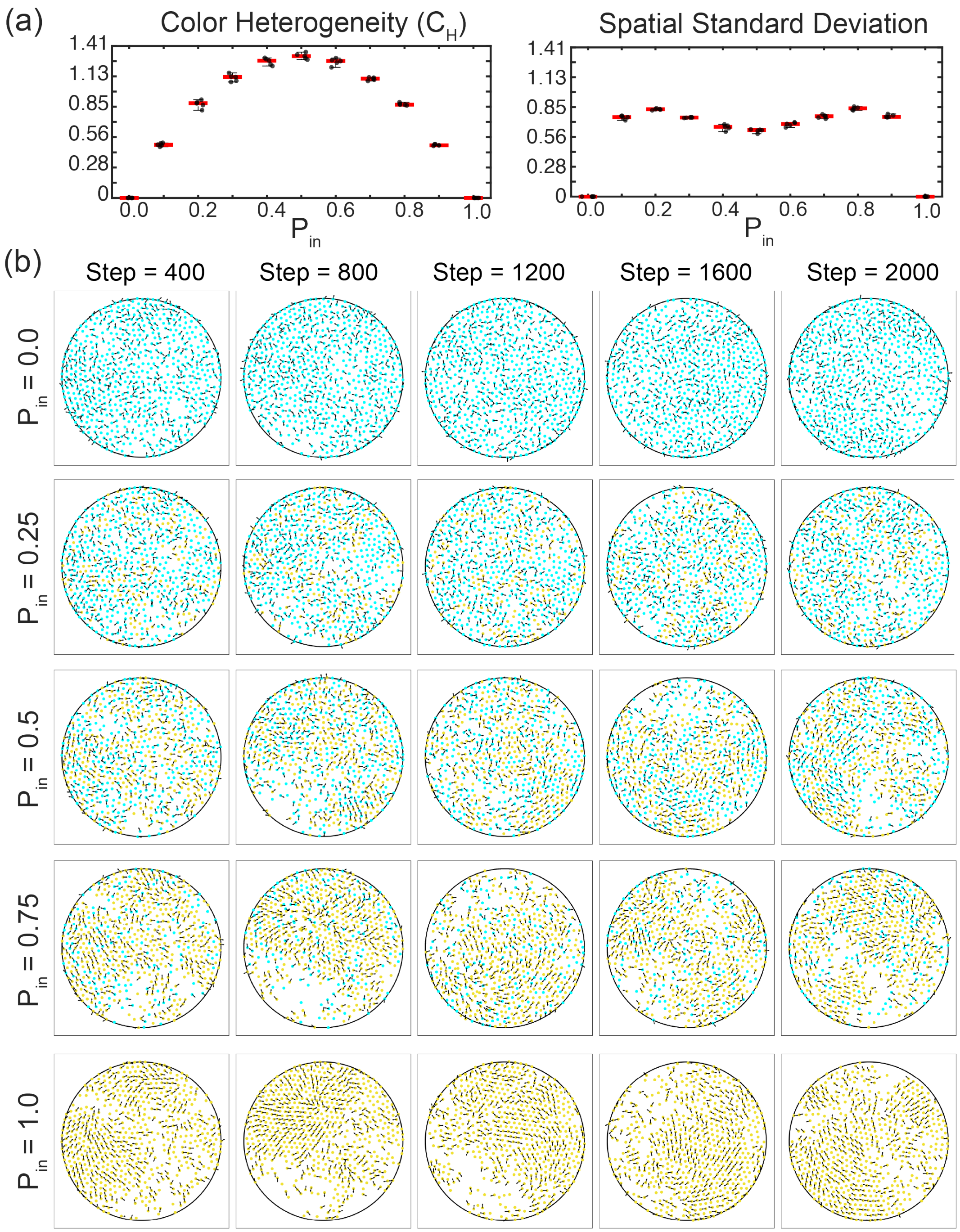


FIG. S8. **Simulation of self-propelled particles with inelastic or elastic interaction rules with different heterogeneity** (a) Local color heterogeneity and its spatial variability as a function of global inelastic particle fraction. Local color heterogeneity was defined as the fraction of neighboring particles (within $r = 3.5$) exhibiting a different collision trait relative to a reference particle. The mean value captures the overall degree of local compositional mixing, whereas the spatial standard deviation characterizes heterogeneity fluctuations across space. Data correspond to independent simulations ($n = 5$) evaluated at the final time point($step = 2000$). (b) Representative simulation snapshots over time ($step = 400, 800, 1200, 1600, 2000$) across the different ratio of inelastic particles.

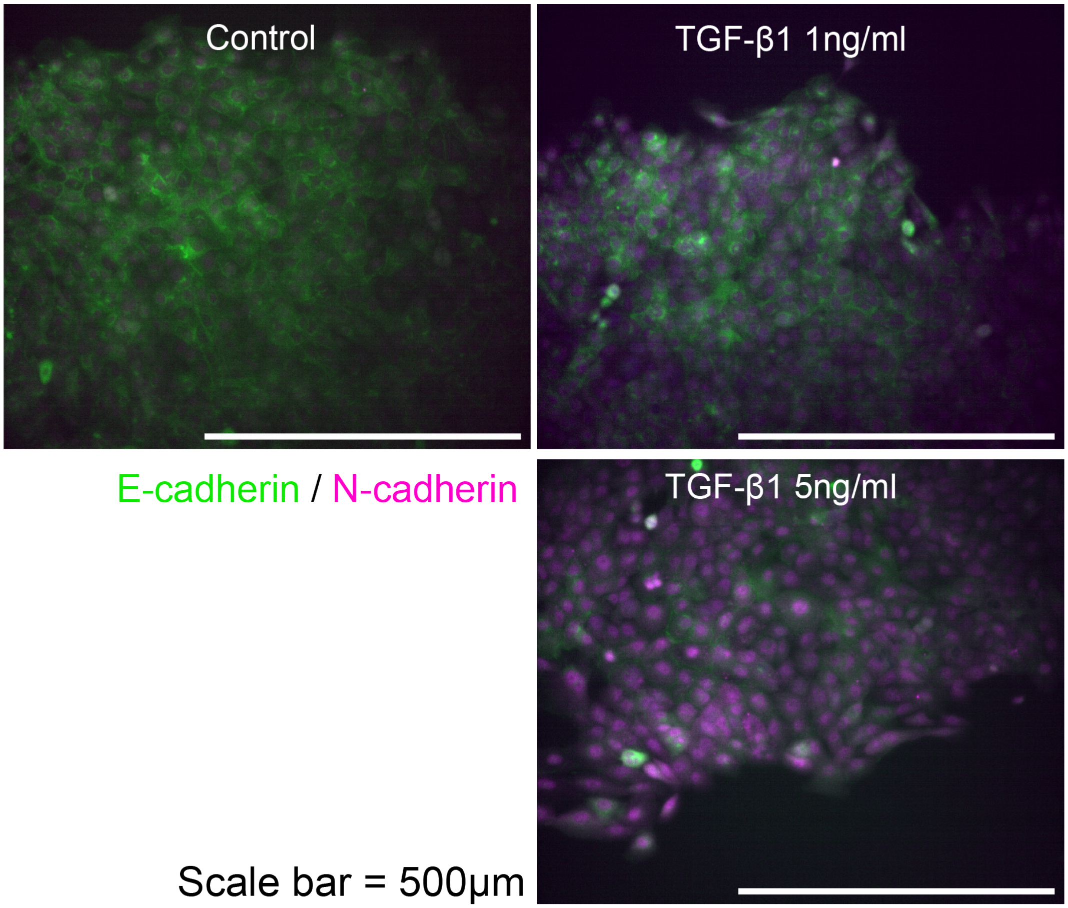


FIG. S9. **Immunofluorescence staining of MCF-10A cells for E-cadherin (green) and N-cadherin (fuchia)** (Top left) Untreated control, (top right) 1 ng/mL TGF-$\beta$1 treatment, (bottom right) 5 ng/mL TGF-$\beta$1 treatment
